# Hippocampal grey matter changes across scales in Alzheimer’s Disease

**DOI:** 10.1101/2025.10.15.682705

**Authors:** Bradley G. Karat, Mahdieh Varvani Farahani, Malcolm Davidson, Arun Thurairajah, Alaa Taha, Taylor W. Schmitz, Ali R. Khan, Alzheimer’s Disease Neuroimaging Initiative

## Abstract

Alzheimer’s disease (AD) is a progressive and debilitating neurodegenerative disease of the central nervous system, characterized by deterioration in cognitive function including extensive memory impairment. The hippocampus, a medial temporal lobe region, is a key orchestrator in the encoding and retrieval of memory and is believed to be one of the first regions to deteriorate in AD. In this work we examined hippocampal macrostructure (specifically gyrification and thickness) and microstructure in Alzheimer’s disease (AD) and mild cognitive impairment (MCI) relative to healthy aged controls in the Alzheimer’s Disease Neuroimaging Initiative (ADNI) dataset. We first utilized an iterative training paradigm to adapt an existing deep learning approach for capturing hippocampal topology to elderly individuals as well as individuals with potential hippocampal degeneration. Using this new model, we found notable decreases in both thickness and gyrification in AD and MCI across both the subfields and anterior-posterior axis. Using the diffusion tensor representation derived from diffusion MRI data, we found significant increases in the mean diffusivity across the extent of the hippocampus in AD and MCI, which may be related to a number of changes such as loss of neuronal cells, decreased fiber density, demyelination, and increased presence of CSF. Examining the primary direction of diffusion relative to canonical hippocampal axes, we found distinct diffusion orientation shifts in AD and MCI throughout the anterior-posterior extent of the subiculum and CA1. Specifically, we found a decrease in diffusion oriented tangentially, and an increase in diffusion oriented along the long-axis. This could potentially be related to the known degeneration of the perforant path, which is greatly affected in AD and is a largely tangential oriented pathway. The AD-related changes in diffusion orientations were found to not have significant spatial overlap with AD-related changes in mean diffusivity, suggesting that they may be capturing distinct spatially-localized disease processes. Finally, we showed that the macro- and microstructure of the hippocampus in AD changed less across age relative to MCI and controls. As well, the age-related hippocampal macrostructure changes in MCI appeared indistinguishable from healthy aging.

## 1. Introduction

Alzheimer’s disease (AD) is a progressive and debilitating neurodegenerative disease of the central nervous system. It is characterized by deterioration in cognitive function including extensive memory impairment, with deficits in episodic memory being one of the first clinical signs of AD (Jacobs et al., 1995; Lane et al., 2017; Masters et al., 2015). The hippocampus, a medial temporal lobe region, is a key orchestrator in the encoding and retrieval of memory and is believed to be one of the first regions to deteriorate in AD (Braak & Braak, 1991; Lisman et al., 2017; Scoville & Milner, 1957; Squire, 2010). The neuropathological changes of AD include the presence of neurofibrillary tangles, accumulation of extracellular amyloid-beta, astrogliosis and microglial activation, and synaptic and neuronal loss (Braak & Braak, 1991; Lane et al., 2017; Serrano-Pozo et al., 2011). The presence of neuronal loss is particularly stark in the entorhinal cortex (ETC) which contains the cell bodies whose axons constitute the perforant path, a major afferent of the hippocampus that is greatly affected in the early stages of AD (Braak et al., 2006; Gómez-Isla et al., 1996; Hyman et al., 1986; Salat et al., 2010). Furthermore, the effect of AD on the hippocampus is not homogenous (Adler et al., 2018). It has been shown that the cells of the Cornu Ammonis (CA) 1 and the subiculum are heavily affected, while the dentate gyrus (DG) and CA3 are relatively spared (Braak et al., 2006; Thal et al., 2002; West et al., 1994). Given its critical role in memory and its early deterioration in AD, the hippocampus has been a prime target for research including biomarker development, prognosis prediction, and potential disease-slowing mechanisms.

Foundational knowledge on the effects of AD on the structure of the brain, including the development of diagnostic criteria, has come from histological examination of neural tissue (Shih et al., 2023). However, there is a need for sensitive and non-invasive probes of neural structure to understand the trajectory of disease development and to characterize disease status in living humans. Diffusion MRI (dMRI) has emerged as a popular technique to measure such neural structure non-invasively, as it is sensitive to the micrometre diffusion of water. On this scale, dMRI can be said to probe microstructure which consists of properties such as cell bodies, axons, dendrites, glial processes, and more. Indeed, dMRI has been used extensively to investigate microstructural changes in mild cognitive impairment (MCI - a risk factor for later developing AD) and AD. Using Diffusion Tensor Imaging (DTI; Basser et al., 1994), studies have generally found a widespread decrease in the fractional anisotropy (FA - quantifies the degree with which diffusion is preferentially oriented along a direction) and an increase in the mean diffusivity (MD - averaged diffusivity) in both white matter and cortical regions (Acosta-Cabronero et al., 2012; Naggara et al., 2006; Spotorno et al., 2023; Stebbins & Murphy, 2009; Weston et al., 2015; Zhang et al., 2009). Applied specifically to the hippocampus, research has shown significant correlations between anterior intrahippocampal MD and memory performance (Yakushev et al., 2010) and glucose metabolism (Fellgiebel & Yakushev, 2011) in patients with early AD. Similar to other cortical regions, a decreased FA and increased MD in the hippocampal GM has been shown in AD compared to age-matched controls (Takahashi et al., 2024; Tang et al., 2016). Diffusion changes have also been noted in the extra-hippocampal WM, including the perforant path, hippocampus-cingulum, and hippocampus-amygdala connections in AD (Chen et al., 2022; Salat et al., 2010). Thus, dMRI appears to be a sensitive technique to investigate the structural changes of the brain and hippocampus non-invasively in AD.

An often underutilized aspect of dMRI is the rich diffusion-orientation information it can provide. Such orientation information forms the basis for diffusion tractography, which attempts to reconstruct the structural pathways of the brain (Jeurissen et al., 2019). The structural pathways of the hippocampus resemble that of the neocortex, with notable radial and tangential components (Ding and Van Hoesen, 2015; Dolorfo & Amaral, 1998; Karat et al., 2024a; Karat et al., 2023; Zeineh et al., 2017). However, the convoluted shape of the hippocampus is reflected in the complexity of its microstructural components, which are oriented relative to this curvature. Previous work has tried to capture such intrahippocampal circuitry in both AD and aged individuals. Shih et al. (2023) found lower tract density connecting the subregions of the hippocampus in AD compared to controls, suggesting potential connectivity loss and hippocampal network disorganization, which are likely related to the cognitive deficits seen in AD (Gelman et al., 2020). Yassa et al. (2010) reported evidence of age-related perforant path deterioration in humans *in vivo* by comparing the primary orientation of diffusion (V1) derived from DTI and *a priori* knowledge of the orientation of the perforant path. Such studies provide valuable information pertaining to the intrahippocampal circuitry, which projects within and across subfields and the hippocampal long-axis. However, many results, including the use of the scalars derived from dMRI as mentioned above (FA and MD), are usually averaged across the whole hippocampus or subfields which inherently averages across many of the intrahippocampal circuit components. As well, it is difficult to compare the tortuous projections provided by tractography between individuals or groups. Thus, no study has examined diffusion orientations granularly across the full extent of the hippocampal grey matter in AD.

In this work, we characterized the macro- and microstructure across the extent of the hippocampus in AD and MCI relative to healthy aged controls. Using HippUnfold, a tool for subfield segmentation and unfolding, we derived macrostructural measures of volume, gyrification, and thickness in a sample of 79 AD, 232 MCI, and 422 controls. We also used DTI to derive measures of MD and V1, where we examined if the orientation of V1 relative to three canonical hippocampal axes (anterior-posterior, proximal-distal, inner-outer) was altered in AD and MCI. We then evaluated if the macro- and microstructural measures were capturing similar changes related to the disease process. Finally, we show how the hippocampal macro- and microstructure change across age within each group, and if these age-related changes are similar between groups. Overall, this work looked to provide a detailed characterization of the hippocampus in AD and MCI, with a focus on a new measure related to the orientation of diffusion.

## 2. Methods

### 2.1 Participants, data acquisition and preprocessing

Data used in the preparation of this article was obtained from the Alzheimer’s Disease Neuroimaging Initiative (ADNI) database (adni.loni.usc.edu). The ADNI was launched in 2003 as a public-private partnership, led by Principal Investigator Michael W. Weiner, MD. The original goal of ADNI was to test whether serial magnetic resonance imaging (MRI), positron emission tomography (PET), other biological markers, and clinical and neuropsychological assessment can be combined to measure the progression of mild cognitive impairment (MCI) and early Alzheimer’s disease (AD). The current goals include validating biomarkers for clinical trials, improving the generalizability of ADNI data by increasing diversity in the participant cohort, and to provide data concerning the diagnosis and progression of Alzheimer’s disease to the scientific community. For up-to-date information, see adni.loni.usc.edu.

In the current study we sourced data from ADNI1, ADNI2, ADNI3, and ADNIGO, with the inclusion criteria that a participant had both T1w and dMRI data. This included a total of 820 participants with a range of 1 to 5 sessions. However, we only utilize the first session from each participant which included both T1w and dMRI data for this cross-sectional analysis. That is, the session analyzed may not have been that participant’s baseline session (since at baseline they may not have undergone both T1w and dMRI acquisitions). The category labels for diagnosis were defined as follows (see the ADNI General Procedure Manual): The classification of AD was given by: (a) memory complaints, (b) abnormal memory function in the Wechsler test, (c) a MMSE score at or below 24, and (d) a dementia rating (CDR) of 0.5 or greater. CN subjects were (a) free of memory complaints, (b) showed normal memory function in the Logical Memory II subscale from the Wechsler Memory Scale, (c) had an MMSE score at or above 24, and (d) had a clinical dementia rating (CDR) of 0. Participants classified within the MCI groups ranged between these extremes. After quality controlling the images, registrations, and hippocampus segmentations, the final number of participants included in the analysis at their respective session dates were 79 with AD (mean (SD) age=76.4 (7.7), F/M=36/43), 232 MCI (mean (SD) age=74.6 (7.9), F/M=108/124), and 422 controls (mean (SD) age=72.7 (7.5), F/M=241/181). We also note that we combined the early MCI, MCI, and late MCI diagnoses into one MCI label for analysis. For the rest of the manuscript we will refer to such classifications as groups rather than diagnoses. See *section 2.2* for more information on quality control, manual correction, and additional model training.

The data used here was acquired across a number of sites and over a period of time (i.e. variation from ADNI 1 to ADNI 3), and as such the specific acquisition parameters may vary per participant. Below we highlight approximate acquisition parameters for each iteration of the ADNI dataset. Detailed information on scanning protocol per scanner can be found at https://adni.loni.usc.edu/data-samples/adni-data/neuroimaging/mri/mri-scanner-protocols/. Briefly, the T1w data from ADNI 1 was acquired on a handful of 1.5T and 3T scanners with an MPRAGE sequence with an approximate TR=2400 ms, minimum full TE, approximate TI=1000 ms, with a voxel size of 1 x 1 x 1.2 mm (Jack et al., 2008). The T1w data from ADNI 2 was acquired on a handful of 3T scanners with an MPRAGE sequence with an approximate TR=2300 ms, minimum full TE, approximate TI=900 ms, with a voxel size of 1 x 1 x 1.2 mm. Finally, the T1w data from ADNI 3 was acquired on a handful of 3T scanners with an MPRAGE sequence with an approximate TR=2300 ms, minimum full TE, approximate TI=900 ms, with a voxel size of 1 mm^3^. In the current study we included 79 participants from ADNI 1/ADNIGO, 195 from ADNI 2, and 459 from ADNI 3.

The standard dMRI acquisition was a SE-EPI sequence with TE=56 ms and TR=7200 ms, with b-values of 0 (typically 5, interleaved) and 1 (typically 46 directions) ms/μm^2^ at a voxel size of 2 mm^3^. As mentioned above, the details of each participant’s acquisition could vary. To ensure more accurate DTI fitting, we excluded all participants who did not have more than 8 directions at the b=1 ms/μm^2^ shell. The mean number of directions for the b=1 ms/μm^2^ shell for each group was 38.50 (AD), 39.67 (MCI), and 39.46 (CN). The diffusion data was preprocessed using snakedwi (version 0.2.1; Khan et al., 2023a), a preprocessing pipeline leveraging snakebids (Khan et al., 2023b) and snakemake (Mölder et al., 2021). Briefly, the snakedwi pipeline used here included denoising (Veraart et al., 2016; Tournier et al., 2019), motion and eddy current distortion correction (Andersson & Sotiropoulos, 2016), Gibbs ring correction (Kellner et al., 2016; Tournier et al., 2019), bias field correction (Tustison et al., 2010), and finally registration to the T1w image space (Yushkevich et al., 2016). Susceptibility-induced distortion correction could not be performed as opposite phase-encoding data was not available.

To supplement the categorical group labels defined via cognitive testing used in this study, we also analysed CSF measures of phosphorylated tau (ptau) and Aβ_42_ in a subset of 348 participants who had such measures available at the same session as their MRI scan. This included 37 participants in the AD group, 98 participants in the MCI group, and 213 participants in the control group. The CSF measures were obtained through lumbar puncture as described in the ADNI procedures manual (http://www.adni-info.org/). The ptau and Aβ_42_ were measured using immunoassay kit-based reagents using previously characterized antibodies (Olsson et al., 2005). Further protocol information on the CSF measures is available in the ADNI procedures manual.

### 2.2 Segmentation, surface modelling, and gradient field generation

To segment the subfields of the hippocampus and to generate surfaces along the hippocampal midthickness surface, we used an automated tool called HippUnfold (version 1.4.1; DeKraker et al., 2022). Briefly, HippUnfold utilizes a deep neural network (a “nnU-net”; Isensee et al., 2021) to provide subject-specific segmentations of the hippocampal grey matter and its topological boundaries along orthogonal directions. The boundary regions include the medial temporal lobe cortex, hippocampal-amygdala transition area, pial surface, stratum radiatum lacunosum moleculare (SRLM), and the indusium griseum. Using these boundaries and the grey matter as the domain of interest, a subject-specific coordinate system is generated by solving Laplace’s equation along the anterior-posterior, proximal-distal, and inner-outer axes of the hippocampus. These coordinates have been shown to have topological correspondence between variably shaped hippocampi (DeKraker et al., 2023), allowing for the projection of subfields defined via histology into the space of each subject. In addition to providing subfield segmentations, HippUnfold also generates surfaces at varying laminar depths across the hippocampal grey matter in both a native and unfolded space. The subfields used in the current work include the subiculum, CA1-3, and CA4 plus the DG which we averaged together here given the resolution. For more technical details refer to DeKraker et al. (2022).

Taking the partial derivative of the Laplace coordinates (a scalar function) with respect to the x, y, and z dimensions provides a gradient field (a vector function), where the vectors at each voxel within the hippocampus point along the anterior-posterior (AP), proximal-distal (PD), or inner-outer (IO) axes (Karat et al., 2023):

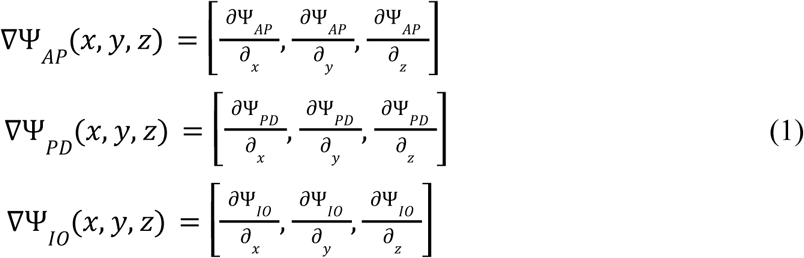

where the function ψ represents the spatial Laplacian coordinates along a particular hippocampal axis (AP, PD, or IO), which were calculated by solving Laplace’s equation ∇^2^ (ψ) = 0) (DeKraker et al., 2022; Karat et al., 2023). These gradients can then be used to capture the orientation of diffusion at each point in the hippocampus, as described in *section 2.4*.

The current nnU-net architectures provided in HippUnfold were trained on healthy adult human data, and may thus not generalize well to the current dataset which includes aged individuals with potential hippocampal degeneration. To improve the nnU-net model for the ADNI dataset, we performed an iterative training paradigm which utilizes successful segmentations from the original nnU-net model. First, HippUnfold was run on the T1w images from all 820 participants using the original T1w nnU-net model. In this first iteration, hippocampal nnU-net segmentations were reviewed by BGK, MVF, MD, AT, and AT and were classified as success (no manual correction or can be improved with manual correction) or failure (cannot be improved with manual correction). If the nnU-net segmentation was successful, it was given a score of 1 to 5 ranging from needing lots of manual correction to no manual correction required. In the first pass of the original model, 1021 left hemisphere nnU-net segmentations were quality controlled, where 1001 succeeded and 20 failed, with the 1001 successful segmentations having scores of 1 (5), 2 (17), 3 (96), 4 (531), and 5 (353). The successful left hemisphere segmentations with a score of 5 (353 segmentations from 198 subjects) were then used to train a nnU-net from scratch. During re-training, a number of right hemisphere segmentations of the original model were quality controlled, where 1006 succeeded and 11 failed out of 1017 reviewed segmentations, with the 1006 successful segmentations having scores of 1 (14), 2 (20), 3 (144), 4 (495), and 5 (339). HippUnfold was then re-ran across all subjects using the newly trained nnU-net. The nnU-net segmentations derived from the new model were then quality controlled based on the following criteria: if a segmentation was deemed a failure or had a score of 3 or less from the first round of QC or was not examined in the first round of QC. In total, 422 left hemisphere updated nnU-net segmentations were quality controlled, where 403 succeeded and 19 failed, with the 403 successful segmentations having scores of 1 (18), 2 (24), 3 (122), 4 (163), and 5 (76).

Out of these left hemisphere segmentations, 81 were manually corrected by BGK for various errors such as oversegmentation of the grey matter label into the CSF or missing anterior hippocampal grey matter, while 42 were not manually corrected and were thus excluded. 231 right hemisphere updated nnU-net segmentations were quality controlled, where all 231 succeeded with scores of 1 (9), 2 (26), 3 (152), 4 (40), and 5 (4). Out of these right hemisphere segmentations, 91 were manually corrected by BGK while 8 were not manually corrected and were thus excluded. All the successful and manually corrected segmentations for the first session of each participant (780 participants in total) were then run through HippUnfold. After quality controlling the diffusion data (registration, quality of the DTI fitting, etc…), a total of 733 participants were included in the final analysis. Supplementary figure 1 shows some examples of the HippUnfold segmentations with the original and re-trained nnU-net models.

### 2.3 Diffusion Tensor Imaging

The FMRIB Software Library (FSL; version 6.0.3, Smith et al., 2004) was used to fit the diffusion tensor using all b = 0 and b = 1 ms/µm^2^ data. DTI characterizes the diffusion process as a symmetric 3×3 tensor where the diffusion signal can be written as:

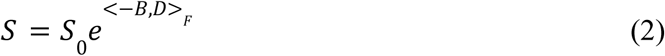

where S is the diffusion signal along a particular direction, B is the b-matrix (Mattiello et al., 1997), D is the diffusion tensor, and <*>_F_ is the Frobenius inner product (Karat et al., 2024a). The eigendecomposition of D provides useful metrics which capture ensemble diffusion characteristics, including three orthogonal vectors V1, V2, and V3 (capturing the orientation of the diffusion process/orientation of the tensor axes), and the mean diffusivity (MD; the mean of the eigenvalues - average diffusivity in physical units). MD was sampled along the midthickness surface (middle of the hippocampal grey matter) for both left and right hemispheres for each participant.

### 2.4 Diffusion relative to the hippocampal axes

As mentioned previously, the orientation of the intrahippocampal microstructure is generally aligned relative to the complex shape of the hippocampus, including its long-axis (anterior-posterior), tangential (proximal-distal; crosses subfields), and radial (inner-outer; crosses laminae) axes (Ding and Van Hoesen, 2015; Dolorfo & Amaral, 1998; Karat et al., 2024a; Karat et al., 2023; Nieuwenhuys et al., 2008; Zeineh et al., 2017). The orientation of diffusion was quantified using the gradient (i.e. vector fields) obtained along the three hippocampal axes (as described in *section 2.2* and equation 1) and the primary orientation of the diffusion tensor (V1; *section 2.3*). Specifically, we calculated scalar cosine similarities between the generated vectors along the AP, PD, and IO axes and V1 at each voxel (figure 1). All vectors were normalized before calculating cosine similarities. The cosine similarity was defined as the inner product between vectors:

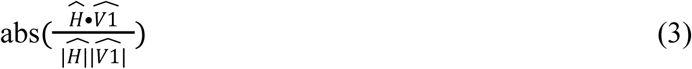

**Figure 1.**
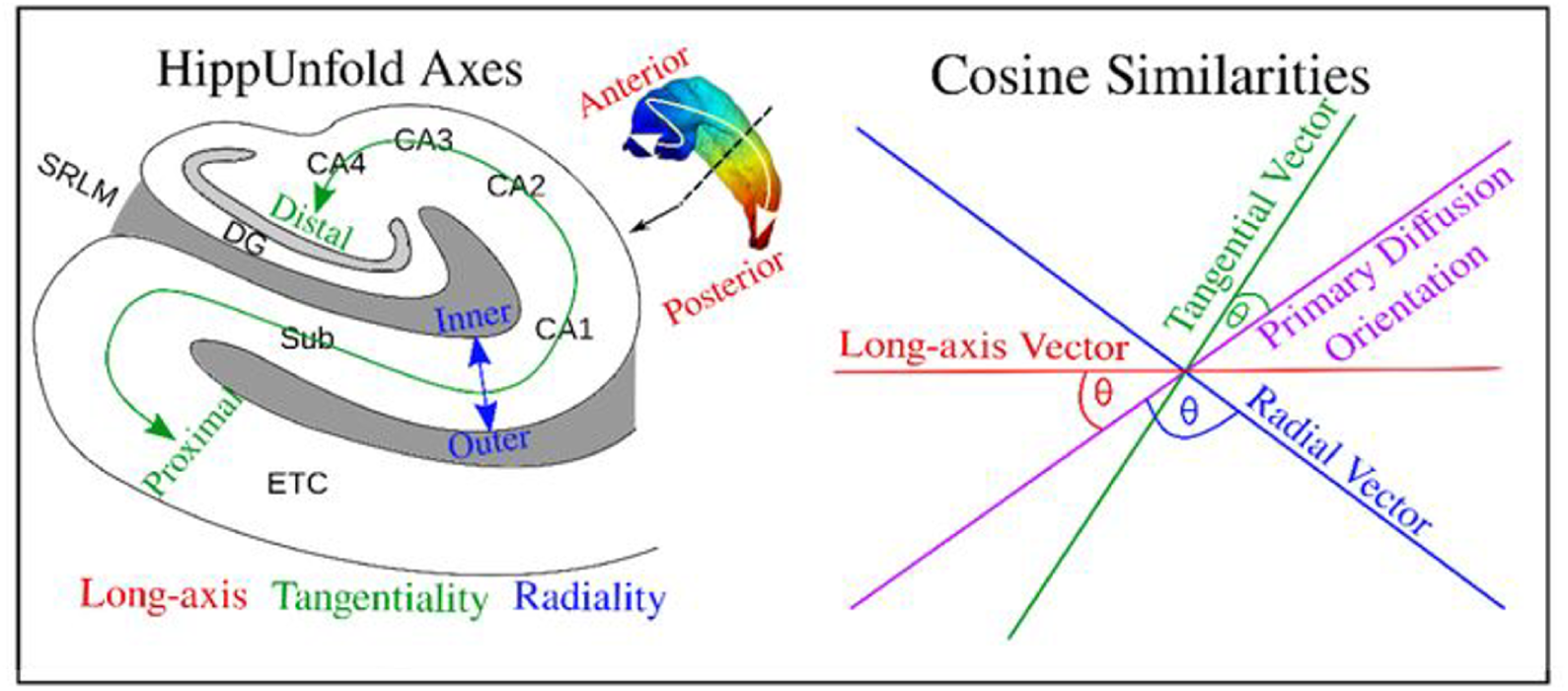
Depicting the calculation of the cosine similarity between the three hippocampal axes (red, blue, and green), and the primary diffusion orientation, quantified in the current study as V1 from the diffusion tensor. Figure adapted from Karat et al., 2023 and Karat et al., 2024a.

where 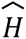 was a hippocampal axis vector in a single voxel (oriented along AP, PD, or IO). The scalar cosine similarity captures the orientation of diffusion at each point within the hippocampus, permitting the investigation of orientation using standard analysis techniques. A cosine similarity of 0 corresponds to a case where diffusion is oriented perpendicular to a particular hippocampal axis, while a cosine similarity of 1 corresponds to a case where diffusion is parallel to a particular hippocampal axis. A total of three scalar cosine similarity metrics were generated for each participant and hemisphere, referred to as the long-axis (AP), tangentiality (PD), and radiality (IO) of diffusion. The cosine similarities were sampled along the hippocampal midthickness surface.

### 2.5 Statistics, correlation, and analysis

A variety of statistics at the subfield-averaged, anterior-posterior averaged, and vertex-wise level were used.

First, linear modelling was used to test if there was a significant interaction between hemisphere and group for each metric examined (volume, gyrification, thickness, MD, long-axis, tangentiality, and radiality of diffusion). For each metric two linear models were built, with one containing all relevant terms (the full model) and another model with all the same terms except for the term of interest (the reduced model; term of interest was the hemisphere-group interaction). An F-test was then performed between the reduced and full model to test if the variable of interest resulted in a significantly better model (i.e. captured significant variance in the dependent variable (DV)). If the F-test was found to not be significant, we averaged hemispheric data for that metric within each participant. The form of the full and reduced model was:

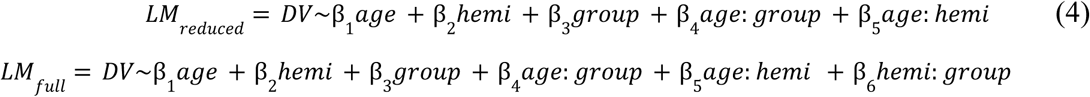

where: implies an interaction between two variables and the DV was a given macro- or microstructural metric. For brevity not all β coefficients were shown here since each interaction with group requires its own coefficients. In this example both models were identical except for an additional interaction term between hemisphere and group. An F-test between the reduced and full model was then used to examine if the hemisphere by group interaction term significantly improved the linear model. A significant F-test suggested that the change in the DV within group was significantly different between hemispheres. False-discovery rate (FDR) correction was applied across all metrics for a given model if any p-value was found to be significant.

Within each subfield-averaged region, Welch’s ANOVA and a subsequent Games-Howell post-hoc test was used to test if there were significant differences in a particular metric across groups. Welch’s ANOVA was chosen given its robustness to unequal variances between groups, and is the preferred choice when sample sizes within each group are unequal, as we have in the current study. Welch’s ANOVA was performed separately for each subfield, and was thus performed 5 times for each metric. We performed false-discovery rate (FDR) correction for the 5 p-values generated from the ANOVA within each metric. Games-Howell post-hoc tests were then performed for each subfield with a significant ANOVA after correction. Thus, 3 p-values (combination of tests between AD, MCI, and controls) were generated for each subfield with a significant Welch’s ANOVA. The Games-Howell post-hoc test inherently provides control of Type 1 errors (Games & Howell, 1976; Sauder & DeMars, 2019). However, we were potentially performing this test multiple times within each subfield. To simplify the analysis, we set a minimum alpha level of 0.01 (assuming we test all 5 subfields/all 5 ANOVAs were significant) for the Games-Howell post-hoc test. The same method described above was repeated for the anterior-posterior parcellations.

To provide increased spatial fidelity, analyses were performed at the surface or vertex-wise level. First, we sought to examine if the age-related changes of a particular metric differed between AD, MCI, and controls. At each vertex on the hippocampal midthickness surface, a linear model was built separately for each group as:

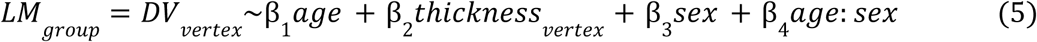

where vertex-wise thickness was included to account for any relation between the DV and the thickness of the hippocampal GM (i.e. to remove effects that may be contributable to partial voluming). The t-statistic for the contrast of age (β_1_) was obtained at each vertex, where the full hippocampal t-statistic map (referred to as age contrast maps; Karat et al., 2024b) depicts linear age-related changes of a given metric within each group. Finally, the age contrast maps were correlated between groups using a hippocampus spin test with 2500 permutations (Karat et al., 2023) to test if the maps have significant spatial overlap. A Bonferroni correction was applied given that for each metric 3 separate spin tests were performed (comparing AD, MCI, and controls). If significantly correlated, then it can be said that the two groups have similar age-related trends of a particular metric across the extent of the hippocampus. No significant correlation would suggest that the age-related trends of a metric are sufficiently different between groups across the hippocampus. Finally, similar models were built as in equation 5, except a contrast of group was included (referred to as group contrast maps), and thus two maps were derived for each metric (the contrast of MCI and controls, as AD is encoded in the intercept). These surface maps highlight which spatial regions of the hippocampus are changing linearly between groups for a particular metric. These maps were then correlated between metrics using the same hippocampus spin test to examine if the macro- and microstructural measures may be capturing similar spatially-specific disease processes. A Bonferroni correction was applied given that for each metric 5 separate spin tests were performed (comparing a particular metric with the 5 other metrics).

## 3. Results

### 3.1 Average macro- and microstructure within group

Figure 2 depicts group-averaged macrostructural measures of gyrification and thickness. Gyrification across AD, MCI, and controls appears to be the largest in the anterior subiculum and CA1 regions, as well as in the DG/CA4, which has been shown previously (DeKraker et al., 2020; Karat et al., 2023). The averaged gyrification appears to be overall lower in the AD group when compared to MCI and controls. Thickness across all groups appears to be highest in the anterior and posterior subiculum, as well as in the DG/CA4 and the anterior CA1. Similar to gyrification, thickness does appear to be lower in the AD group.

**Figure 2.**
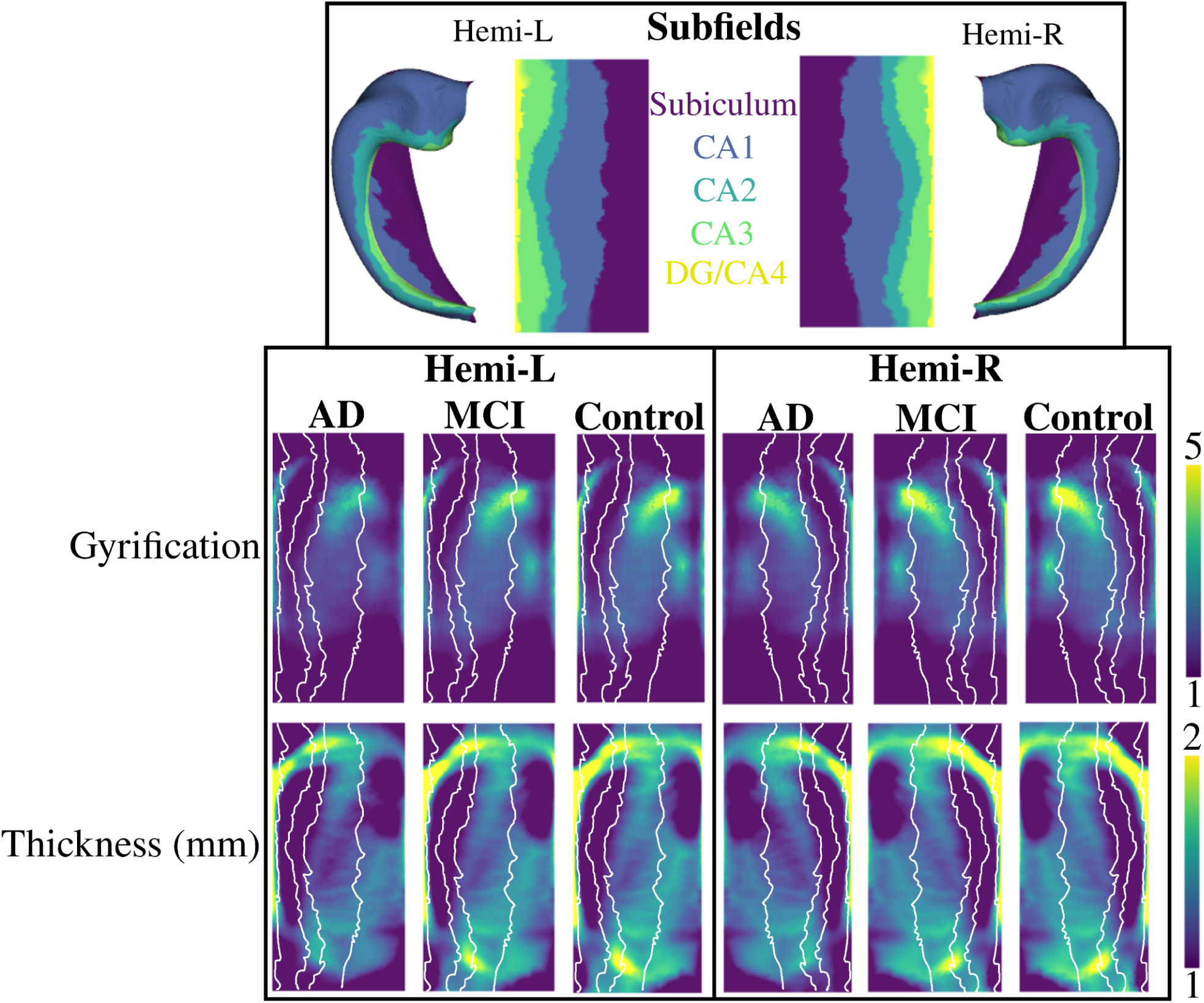
Macrostructural metrics of gyrification and thickness averaged within each group and shown in unfolded space. The top row shows the subfields in both folded and unfolded space. The white lines overlaying the two bottom rows depict the subfield borders. CA - cornu ammonis; DG - dentate gyrus; AD - Alzheimer’s Disease; MCI - Mild Cognitive Impairment.

Figure 3 shows the group-averaged microstructural measures of mean diffusivity (MD), and the long-axis, tangential, and radial orientation of the primary direction of diffusion (the cosine similarity; see *section 2.4*). MD is lowest in CA1 and highest in the subiculum and CA2/CA3, which has been shown previously (Karat et al., 2023). There is a large increase in the averaged MD in the AD group compared to MCI and controls. The primary direction of diffusion is largely oriented along the hippocampal long-axis in the body and posterior subiculum, as well as in the body of the DG/CA4, CA3, and CA2. As well, the long-axis diffusion appears to be increased in the AD group. Diffusion is oriented tangentially (crosses the subfields) mainly in the anterior and body of CA1 and CA2. Diffusion is less tangentially oriented in AD when compared to MCI and controls. There does not appear to be a region of very high radially oriented diffusion, suggesting that a lot of the primary orientation of diffusion is in the long-axis to tangential plane of the hippocampus.

**Figure 3.**
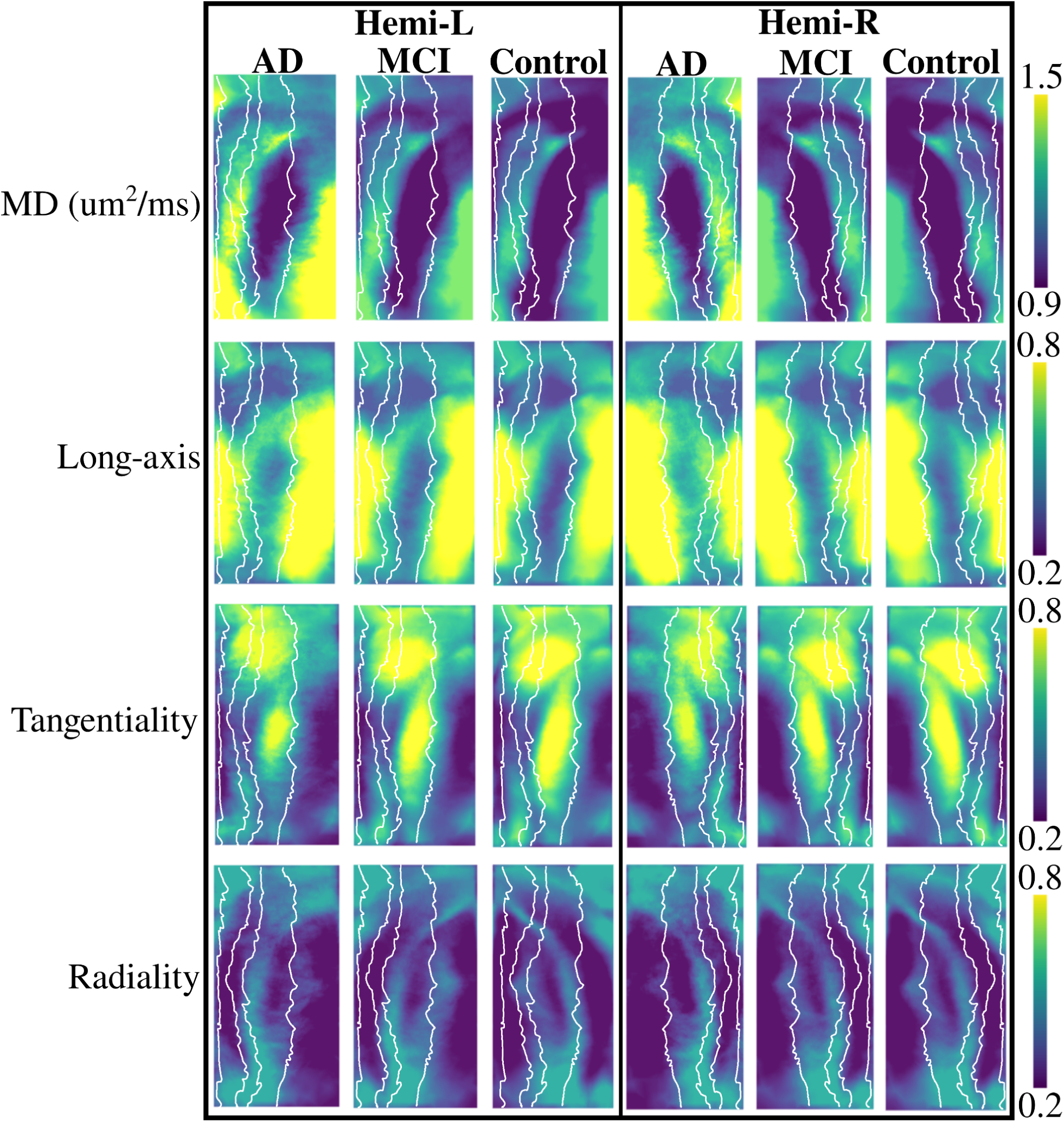
Microstructural metrics of mean diffusivity (MD) and long-axis, tangential, and radial oriented diffusion averaged within each group and shown in unfolded space. The white lines depict the subfield borders, seen in the top of figure 2.

### 3.2 Group-related changes in subfield macro- and microstructure

Subfield-averaged values of macro- and microstructure between AD, MCI, and controls is shown in figure 4. No significant interaction was found between hemisphere and group for volume (F(1,7290)=0.08, p=0.92), thickness (F(1,7290)=0.39, p=0.68), gyrification (F(1,7290)=0.06, p=0.94), MD (F(1,7290)=0.87, p=0.42), long-axis (F(1,7290)=1.86, p=0.16), tangential (F(1,7290)=0.30, p=0.74), or radial (F(1,7290)=0.48, p=0.62) oriented diffusion (supplementary figure 2). Thus, hemisphere data (left and right hemisphere within each participant) was averaged.

**Figure 4.**
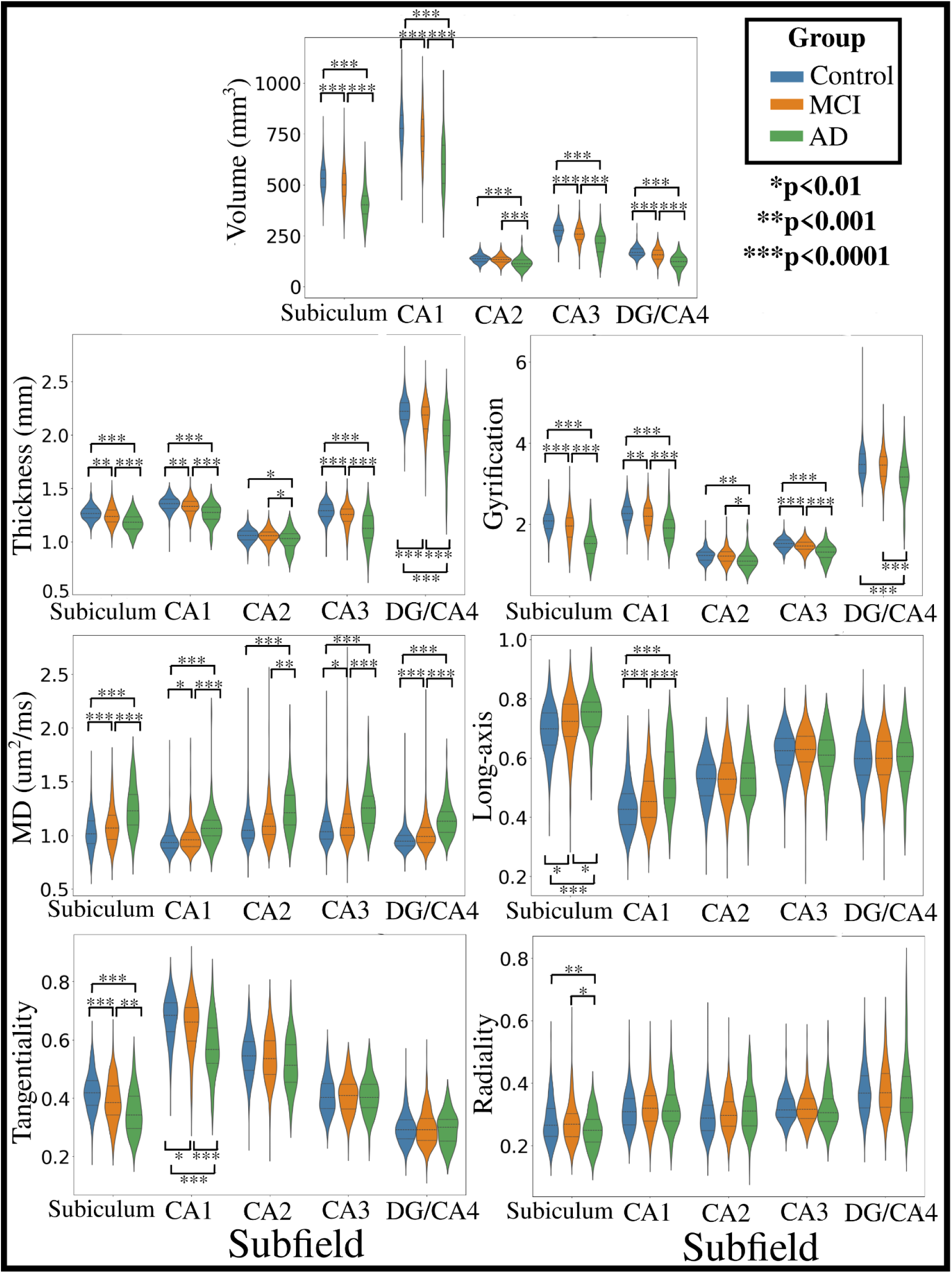
Subfield and hemisphere averaged distributions of macro- and microstructure within each group. Significant differences between groups are represented by asterisks, calculated with a Games-Howell post-hoc test.

The statistics for each comparison within a metric and subfield is presented in supplementary table 1. Widespread differences in subfield macro- and microstructure can be seen between the 3 groups (figure 4). The volume of each subfield is significantly smaller in AD compared to MCI and controls, while MCI has significantly smaller volume compared to controls in all subfields but CA2. The differences in thickness and gyrification are similar to each other, with both being significantly lower (thinner and less gyrification) in AD compared to MCI and controls. The MCI group also has significantly lower thickness than controls in all subfields except CA2, and less gyrification in all subfields except CA2 and DG/CA4. Related to the microstructure, the MD was significantly higher in AD than MCI and controls in all subfields, while MCI had significantly higher MD in the subiculum, CA1, CA3, and DG/CA4 when compared to controls. The changes in the diffusion orientations (long-axis, tangentiality, and radiality) were less global, with changes mainly seen in the subiculum and CA1. Diffusion in the subiculum and CA1 was increasingly oriented along the long-axis in AD when compared to MCI and controls, while there was lower tangential oriented diffusion in these subfields.

### 3.3 Group-related changes in anterior-posterior macro- and microstructure

Similar to *section 3.2*, figure 5 shows values of macro- and microstructure between AD, MCI, and controls averaged across the anterior-posterior axis which is parcellated into head, body, and tail. The statistics for each comparison within a metric and anterior-posterior parcellation is presented in supplementary table 2. The overall trends are similar to that seen in figure 4.

**Figure 5.**
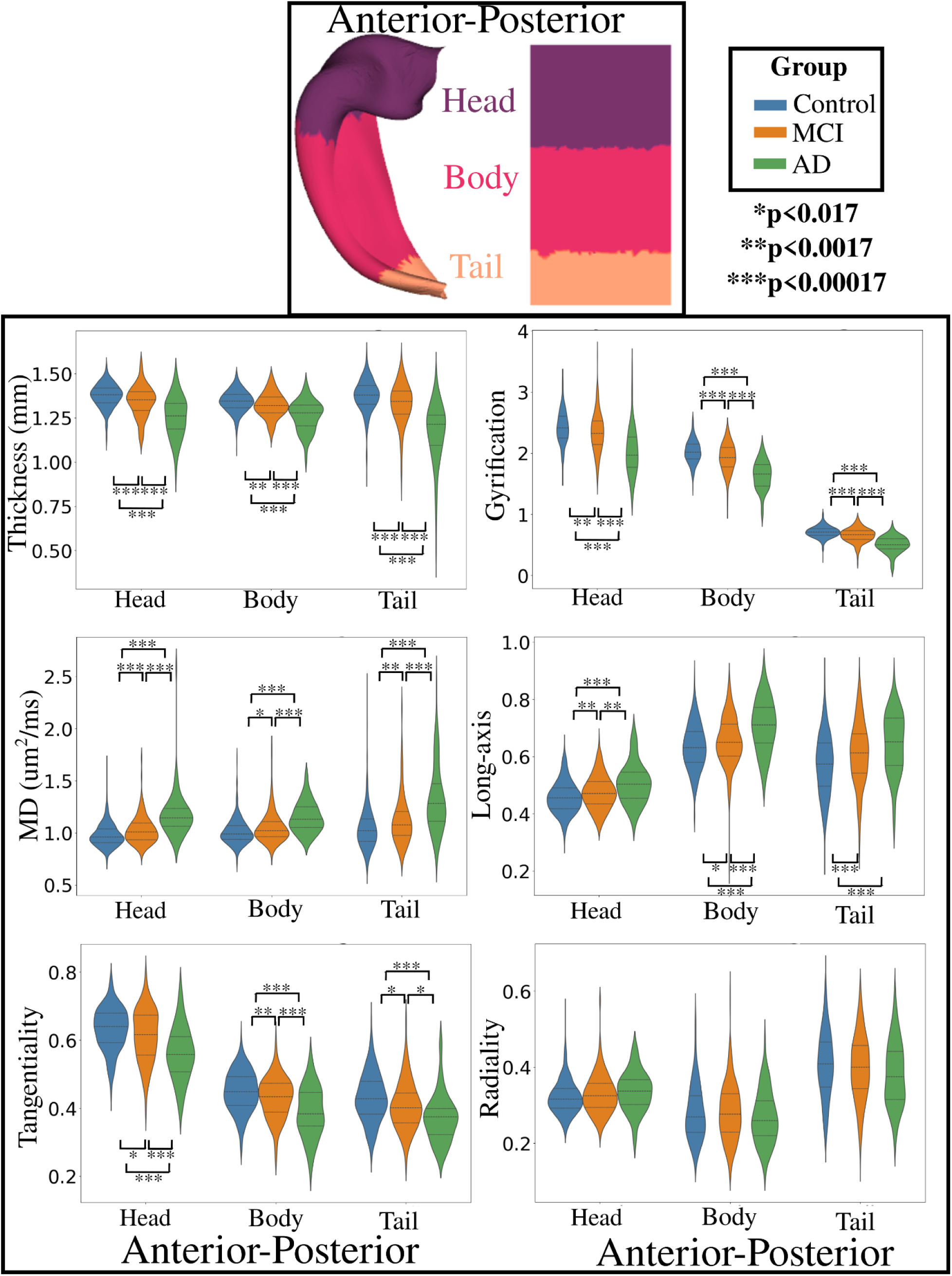
Anterior-posterior and hemisphere averaged distributions of macro- and microstructure within each group. Significant differences between groups are represented by asterisks, calculated with a Games-Howell post-hoc test.

Significant differences in both thickness and gyrification can be seen between AD, MCI, and controls across the head, body, and tail. The mean diffusivity was significantly higher in AD compared to MCI and controls, and between MCI and controls across the anterior-posterior axis. There are also significant diffusion orientation shifts, with higher long-axis oriented diffusion and lower tangential oriented diffusion across the head and body between AD, MCI, and controls. Overall, there are appreciable differences in the macro- and microstructure of the hippocampal head, body, and tail between AD, MCI, and controls.

### 3.4 Spatially distinct changes in macro- and microstructure across group

Surface t-statistic maps which capture spatial changes in a particular metric across group (referred to as group contrast maps) were generated (top row of figure 6). For the contrast of controls (top left of figure 6), it can be seen that the increase in gyrification and thickness relative to AD (as noted in *sections 3.2* and *3.3*) is ubiquitous across the hippocampus, except for part of the head and body of CA2, where the t-statistic is closer to 0. Similarly, the lower MD in controls relative to AD is seen across the extent of the hippocampus, except for in the body of CA1 and the body and tail of CA2. Sharp and more spatially localized changes in the diffusion orientations relative to the long-axis and the tangential axis is seen, with changes largely occurring in the extent of CA1 and at the border with the subiculum. Interestingly, while less change was noted between controls and AD in the diffusion orientations relative to the radial axis in *sections 3.2* and *3.3*, there is a distinct pattern of relatively large increases and decreases which cuts across both the subfields and the anterior-posterior regions. For example, in the anterior CA1 there is a substantial decrease in the radially-oriented diffusion in controls relative to AD, but in the body and tail of CA1 there appears to be regions of increased radial diffusion. Similar results to those described above is noted for the contrast of MCI, though with an apparent attenuation in the magnitude of the t-statistic. That is, the difference between controls and AD across the extent of the hippocampus are larger than the differences between MCI and AD, though the changes appear to be occurring in the same spatial locations.

**Figure 6.**
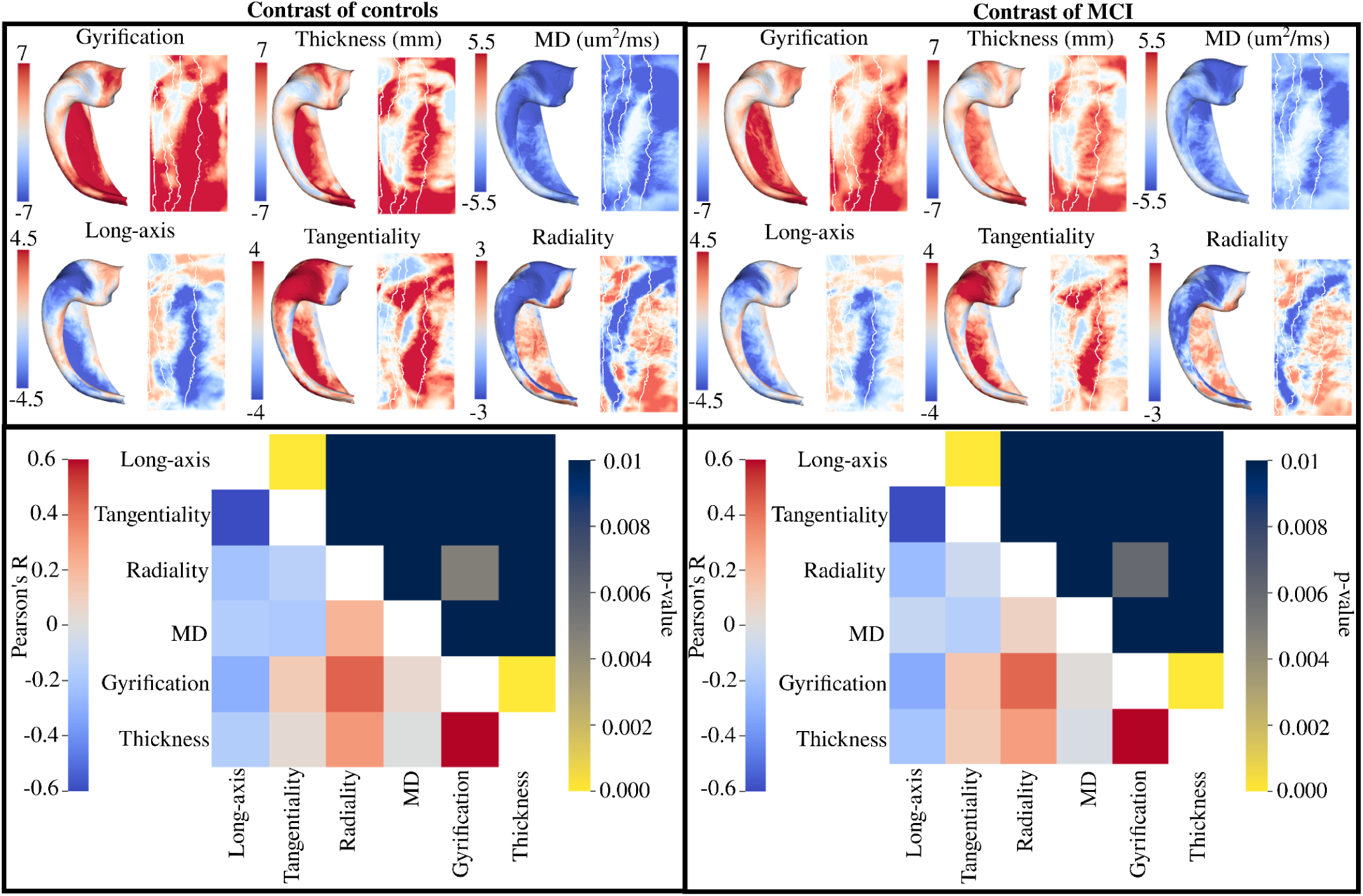
Vertex-wise group contrast t-statistic maps for macro- and microstructure. Maps were calculated with hemisphere-averaged data and plotted on a left hippocampal surface. Bottom row depicts the Pearson’s R correlation and p-values derived from a spin test between the group contrast maps for all metrics. Note that the colour bar for the p-values is inverted such that any brighter component of the upper triangle of the heatmap corresponds to a significant p-value (p<0.01).

The bottom row of figure 6 depicts the correlation between all of the t-statistic maps in the top row using a hippocampus spin test (Karat et al., 2023). The correlation describes the extent of the spatial overlap between any two of the t-statistic maps. Thus it is testing if two metrics are capturing similar differences between groups across the extent of the hippocampus. If the correlation is significant then that suggests that those two metrics are capturing similar spatial changes between groups, while if not significant that suggests that the changes of the two metrics between groups are spatially distinct. For the contrast of controls, the correlation between the long-axis and tangential oriented diffusion (R=-0.7, p<0.01), gyrification and radial oriented diffusion (R=0.44, p<0.01), and gyrification and thickness (R=0.62, p<0.01) is significant. This suggests that the macrostructural differences between controls and AD are largely occurring in the same spatial regions, and that the diffusion orientation changes are occurring along the tangential and long-axis. Interestingly, MD was not significantly correlated with any other metric, suggesting that the changes in overall diffusivity may be spatially distinct from the changes between controls and AD in other measures. The correlations for the contrast of MCI were very similar, with significant correlation between the long-axis and tangential oriented diffusion (R=-0.7, p<0.01), gyrification and radial oriented diffusion (R=0.43, p<0.01), and gyrification and thickness (R=0.61, p<0.01).

### 3.5 Age-related changes in macro- and microstructure within group

Similar to *section 3.4*, t-statistic maps with a contrast of age were derived separately for each group. Such age contrast maps capture linear changes in a particular macro- or microstructural metric across age within each group. The age contrast maps were then correlated between the groups using the same spin test as described in *section 2.5*. If the maps are significantly correlated then that would suggest that the two groups have spatially similar changes in a particular metric across age, while if not significant that suggests that the age related trends are sufficiently different across the extent of the hippocampus. In figure 7 the age contrast maps are shown for macrostructure. It can be seen that the age related changes for gyrification and thickness in the AD group are not significantly correlated with the age related changes of the MCI or control group. Opposingly, the age related changes of both macrostructural measures are significantly correlated between MCI and controls. As well, it appears that macrostructure is changing more across age in the MCI and control group (larger overall *t*-values) as compared to the lower *t*-values of the AD group.

**Figure 7.**
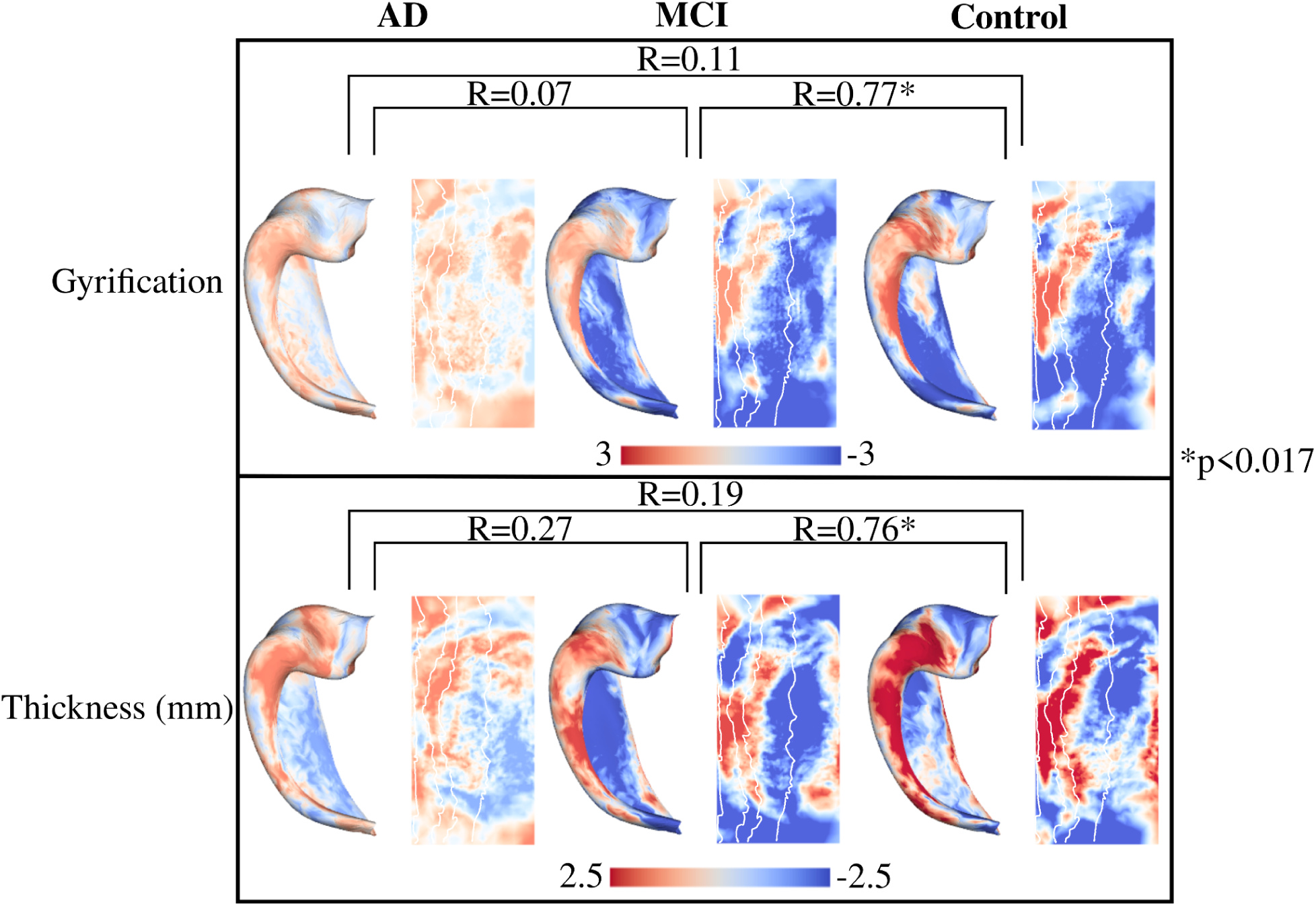
Vertex-wise age contrast maps within group for macrostructure. Maps were calculated with hemisphere-averaged data and plotted on a left hippocampal surface. Spin test correlations were performed for each age-contrast map across groups. The asterisk represents a significant p-value (p<0.017).

In figure 8 the age contrast maps are shown for the microstructure measures. There is no significant correlation between any group for the age related changes of MD. As well, MD appears to increase more across age in the control group when compared to the AD and MCI group (larger overall *t*-values). For the diffusion orientations, the age related changes are significantly correlated between MCI and controls, while there is no significant correlation with AD. Similar to the macrostructure, the diffusion orientations appear to change more across age in the MCI and control groups when compared to the AD group.

**Figure 8.**
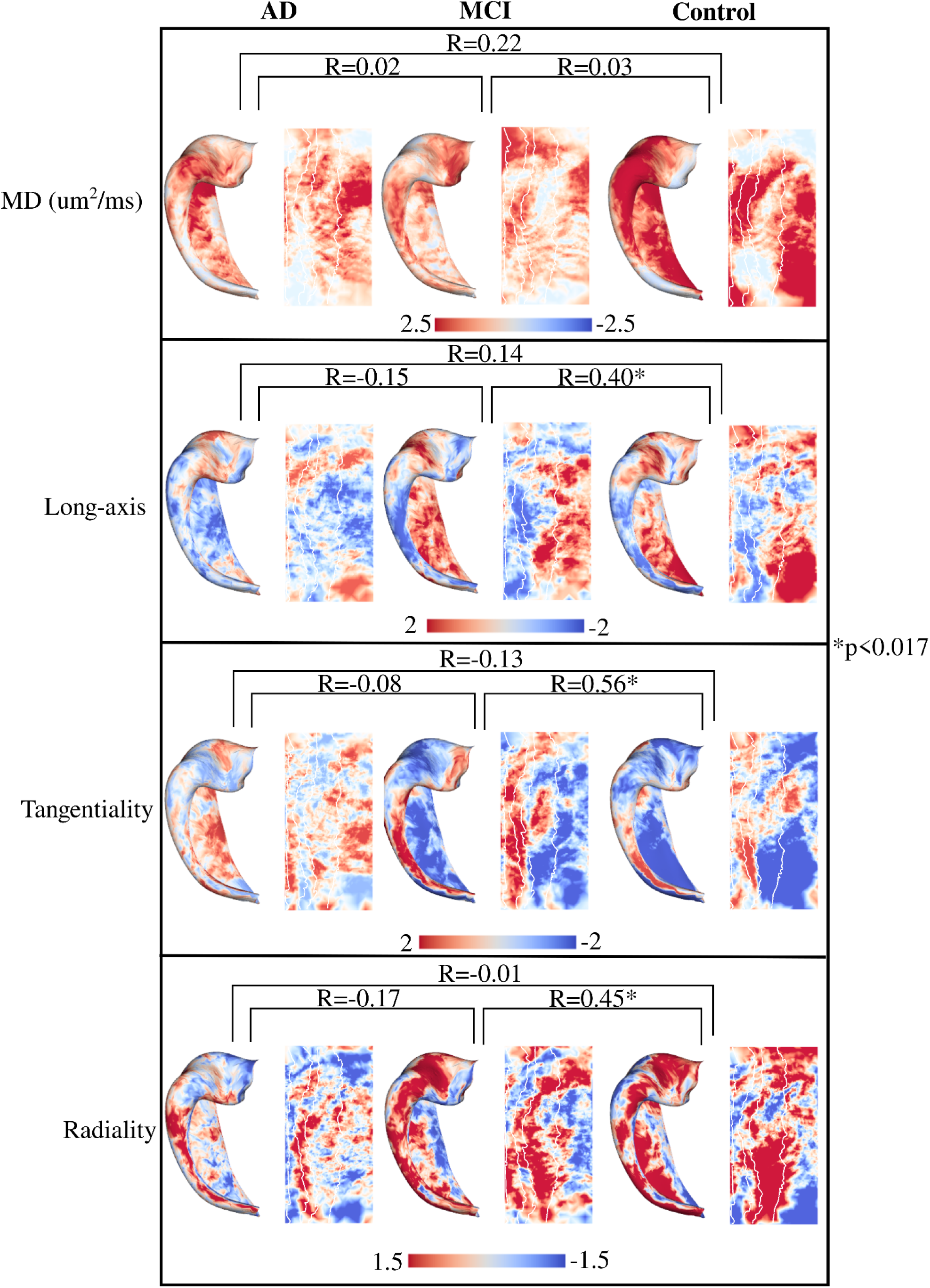
Vertex-wise age contrast t-statistic maps within groups for microstructure. Maps were calculated with hemisphere-averaged data and plotted on a left hippocampal surface. Spin test correlations were performed for each age-contrast map across groups.

## 4. Discussion

### 4.1 Widespread macro- and microstructural changes between AD, MCI, and controls

The hippocampus is a region which is known to deteriorate in AD (Braak & Braak, 1991; Lisman et al., 2017; Squire, 2010). Much of our knowledge on the effects of AD on the hippocampus and other cortical regions has come from histological work examining neural soma and processes (Braak & Braak, 1991; Braak et al., 2006; Shih et al., 2023). However, there has been much contemporary research focused on characterizing AD through a variety of non-invasive measures, typically using MRI. Common measures derived from structural MRI include gyrification and thickness. A variety of cortical regions have been found to be hypo- or hyper-gyrified in AD compared to MCI and controls. Liu et al. (2012) found that global cortical gyrification decreased with increasing severity of AD. Lebed et al. (2013) and Núñez et al. (2020) found that individuals with AD had increased gyrification in the entorhinal cortex and decreased gyrification in the insular cortex when compared to controls. They suggest that such a gyrification pattern occurs because of the dynamic changes induced as a consequence of atrophy in AD, which is prominent in the temporal lobe (Barnes et al., 2009). Indeed, longitudinal work has shown dynamic sequences of cortical atrophy beginning in limbic and temporal areas before moving into higher-order association areas (Thompson et al., 2007). In the current study we found significantly lower gyrification in AD compared to MCI and controls across the extent of the hippocampus, including all the subfields, as well as significantly lower gyrification in MCI compared to controls except in CA2 and DG/CA4. We also found significantly smaller volumes of all subfields in AD compared to MCI and controls, and in MCI compared to controls except for CA2, indicative of widespread hippocampal atrophy. As suggested above, it is likely that extensive hippocampal atrophy would be correlated with changes in gyrification. Similar to the trends in gyrification and commensurate with atrophy in AD is a reduction in the thickness of the grey matter, which has been shown throughout the extent of the cortex (de Miras et al., 2017; Hwang et al., 2016; Singh et al., 2006; Thompson et al., 2007; Querbes et al., 2009; Youn et al., 2021). We found significantly lower thickness in AD compared to MCI and controls across the hippocampus, including all the subfields, as well as significantly lower thickness in MCI compared to controls except in CA2. The hippocampus is important for encoding perception in both spatial and temporal domains, and the extensive atrophy with reduced gyrification and thickness ostensibly related to neuron loss, contribute to the cognitive deficits seen in AD and MCI.

While useful for examining the overall structure of the cortex and hippocampus, the macrostructural measures described above are inherently coarse. Measures derived from diffusion MRI may improve sensitivity to changes in AD and MCI. Indeed, previous research has indicated widespread increases in MD in patients with AD and MCI which precede volume loss, with damage to the white matter appearing to begin in the temporal lobe (Acosta-Cabronero et al., 2012; Naggara et al., 2006; O’Dwyer et al., 2011; Spotorno et al., 2023; Stebbins & Murphy, 2009; Weston et al., 2015; Zhang et al., 2009). A plethora of diffusion changes have been previously shown within the hippocampal grey matter, including increased FA, decreased MD, decreased stick fraction and increased isotropic fraction (Hong et al., 2012; Shahid et al., 2022; Takahashi et al., 2024; Tang et al., 2016; Weston et al., 2015; Yakushev et al., 2010). Such an increase in diffusivity and decreased stick fraction could be related to increased partial voluming (i.e. the presence of CSF), loss of neuronal cells (particularly the pyramidal cells of the hippocampus), demyelination, fiber density, alterations in axon diameter, and/or increased glial response. Diffusivity in the hippocampus has also been correlated with tau burden as a continuous measure of disease pathology, cognitive scores with tasks dependent on hippocampal function, and glucose metabolism in AD (Chen et al., 2022; Hong et al., 2012; Shahid et al., 2022; Yakushev et al., 2010; Fellgiebel & Yakushev, 2011). This suggests that hippocampal diffusion changes are related to disease processes which may convey important information on cognitive status and outcomes in AD. Changes have also been found in the extrahippocampal white matter, including the hippocampal-cingulum and the perforant path, suggesting that the hippocampal afferents and efferents are affected as well (Chen et al., 2022; Salat et al., 2010; Yassa et al., 2010). In the current study we found similar changes, with a large increase in MD across both the subfields and anterior-posterior axis for AD, MCI, and controls. This suggests that hippocampal grey matter diffusion is sensitive to disease processes related to both MCI, which tend to be more subtle compared to controls, and to AD where structural alterations are more extensive.

### 4.2 Diffusion orientation changes in AD and MCI

Diffusion MRI often provides rich orientation information which forms the basis for diffusion tractography, which attempts to reconstruct the structural pathways of the brain (Jeurissen et al., 2019). Diffusion tractography has been used extensively to quantify changes to both specific white matter bundles and global structural connectivity (Feng et al., 2024; Lee et al., 2015; Lo et al., 2010; Reginold et al., 2016; Shaikh et al., 2022; Xie et al., 2005). For example, reduced FA and increased MD of the cingulum and cingulum-angular bundles have been shown in AD, with significant correlations between the diffusivity of the cingulum and hippocampal volumes (Lee et al., 2015; Xie et al., 2005). Similarly, Shaikh et al. (2022) found increased diffusivity and lower tract volume in the fornix of AD patients, which was correlated with hippocampal volume. These results suggest that the efferents of the hippocampus (i.e. the fornix) are significantly affected in AD. Fewer studies have examined the diffusion orientations in the grey matter, largely because of the lower anisotropy, where there may be no primary microstructural orientation. However, the cortical column generally has a structure of distinct radial and tangential components, including the hippocampus whose microstructure is oriented relative to its curved structure (Ding and Van Hoesen, 2015; Dolorfo & Amaral, 1998; Karat et al., 2024a; Karat et al., 2023; Zeineh et al., 2017). Using tractography within the hippocampal grey matter, Shih et al. (2023) found lower tract density connecting the subregions of the hippocampus in AD vs. controls, suggesting potential intrahippocampal network disorganization. In a similar fashion to the current study, Lee et al. (2020) measured the dot product of the normal vector to the cortical surface (i.e. the radial orientation) with the primary diffusion orientation from DTI (V1) in AD. They found decreased radially-oriented diffusion in the entorhinal, insula, frontal, and temporal cortex which correlated with disease progression. They also showed that the radiality measure could delineate between controls and early MCI. However, they did not analyze the hippocampus, likely due to its complex curved organization making it difficult to define its radial orientation. In the current study we calculate a similar measure as in Lee et al. (2020) within the hippocampus by using a bespoke software to fully capture the 3D extent of the hippocampus. We also quantified the additional dimensions of diffusion, namely the tangential and long-axis or anterior-posterior orientations, along with the radial orientation. We found that the orientation changes were largely confined to the subiculum and CA1 across the anterior-posterior extent of the hippocampus. A significant decrease in the tangentiality of diffusion in AD to MCI to controls, and a corresponding significant increase in the long-axis orientation was found in both CA1 and the subiculum. We also found a significant decrease in the radiality only in the subiculum between AD and MCI and controls. Some of the most notable structural changes of the hippocampus in AD are the presence of neurofibrillary tangles and cell loss in the subiculum and CA1 (Braak et al., 2006; Thal et al., 2002; Van Hoesen & Hyman, 1990; West et al., 1994). As well, neuronal loss is extensive in the entorhinal cortex, affecting the cell bodies whose axons form a major afferent of the hippocampus called the perforant path (Braak et al., 2006; Gómez-Isla et al., 1996; Hyman et al., 1986; Salat et al., 2010). The perforant path projects into the subiculum on its way to the granule cells of the dentate gyrus and the apical dendrites of the CA subfields, where a large part of its trajectory appears to be tangential (Dolorfo & Amaral, 1998; Duvernoy et al., 2013; Zeineh et al., 2017). In the current study, the significant decrease in the tangentiality of diffusion particularly in the subiculum and CA1 may correspond to degradation or alterations of the perforant path in MCI and AD. Indeed, we found correlations of MD within a segmentation of the expected spatial location of the perforant path and subfield-averaged tangentiality across all participants (supplementary figure 3). In particular, CA1 tangentiality and perforant path MD were significantly negatively correlated (R=-0.41, p<0.01; supplementary figure 3), suggesting that alterations of the perforant path may be driving some of the changes seen in intrahippocampal diffusion orientations.

Another goal of the current study was to examine if the changes in diffusion orientations align with the changes in MD. That is, are the regions of known MD changes overlapping with the regions of diffusion orientation changes. By examining the vertex-wise spatial changes of each metric relative to group label, we found that the diffusion orientations did not have significant spatial overlap with MD (figure 6). While MD appeared to have larger global changes in AD relative to MCI and controls, the changes in the diffusion orientations were more localized.

Furthermore, the changes in MD in AD relative to MCI and controls was lowest in CA1, while CA1 tended to be the region where the diffusion orientations changed the most (figure 6).

This may suggest that the measures are capturing distinct spatially localized disease processes. We did find significant spatial overlap between thickness and gyrification, suggesting that the macrostructural changes in AD occur in similar spatial regions.

### 4.3 Variable changes in macro- and microstructure across age between AD, MCI, and controls

While the pathogenesis of AD is distinct from aging, it is one of the most critical risk factors as older individuals have an increased incidence of the disease (Herrup, 2010; Kawas et al., 2000; Nelson et al., 2011). Indeed, in the non-familial form of AD (i.e. the sporadic form), it has been posited that some adverse event combined with aging is what triggers the beginning of the degenerative process, eventually leading to AD (Herrup, 2010). It has also been suggested that younger AD patients have significantly faster atrophy rates and cognitive decline (Barnes et al., 2009; Fiford et al., 2018). Considering a wider age range, Holland et al. (2012) found reduced rates of atrophy across age for AD and MCI (changes less across age), whereas healthy controls showed increased rates of atrophy across age, presumably associated with the normal aging process. Partly commensurate with the current study, we found that the macro- and microstructure of the hippocampus in AD appeared to change less across age compared to MCI and controls (lower absolute *t*-values in figure 7 and 8) across a relatively wide age range of 55.3 years to 93.7 years. This may suggest that once the degenerative process of AD has begun that aging becomes a less impactful feature for the structural degeneration of the hippocampus. It is also likely that the changes of the hippocampus across healthy aging are subtle relative to the considerable atrophy of the hippocampus across AD progression. We also found that spatially the age-related changes of macro- and microstructure in AD were not correlated to the spatial changes in MCI and controls, suggesting potentially different mechanisms of change across age in AD. We did find significant correlations of the age-related changes between MCI and controls in all metrics except for MD. In particular, the similarity in the age-related changes for gyrification and thickness between MCI and controls is quite conspicuous, suggesting that relative to hippocampal macrostructure aging in MCI is similar and perhaps not distinguishable from healthy aging. An important note however is that the current study was cross-sectional, and future analyses with longitudinal data will be needed to investigate this further.

### 4.4 Limitations

One limitation of the current study was the use of cognitive scores for grouping AD. Definitive diagnosis of AD requires cognitive impairment with a memory component, along with substantial numbers of neurofibrillary tangles and amyloid plaques confirmed at autopsy (Braak et al., 1993; Mirra et al., 1991; Nelson et al., 2011). Thus there is a risk that some participants may be misdiagnosed, potentially confounding the current group-based analysis. To supplement our analysis, we analyzed CSF measures of ptau and Aβ_42_. Such CSF measures have been of recent interest, given that CSF circulates within and around the brain and thus potentially carries sensitive information on disease status. Previous work has shown good correspondence between tau and amyloid-β in both PET-classified and autopsy-confirmed AD (Hansson et al., 2019; Shaw et al., 2009). In supplementary figure 4 we show that the ratio of ptau and Aβ_42_ was significantly different between the group labels used in the current study. In supplementary figure 5 we show the correlation of the subfield-averaged measures used in the current study with the ratio of ptau and Aβ_42_. Thus while the use of cognitive scores for subject grouping remains a limitation of the current study, the difference of such CSF measures between the groups may increase our confidence in such labels.

Another limitation of the current study was the necessary exclusion of highly degenerated hippocampi, given the difficulty in getting a quality segmentation and surface reconstruction in such cases. This means that our analysis is missing some cases of highest AD severity, though one might expect that their inclusion would make the differences between AD and MCI and controls more conspicuous. Another limitation was the 2mm^3^ resolution of the diffusion data. Given the thickness of the hippocampus, at such resolutions there are likely partial volume effects (PVE) with surrounding white matter and CSF. As well, given the decrease in thickness in both the AD and MCI groups, it is likely that such partial voluming may be significantly different between the groups. However, we did attempt to minimize such PVE by sampling all measures along the middle surface of the hippocampal gray matter and by adding thickness as a covariate in the linear models built on the hippocampal surfaces (*sections 3.4* and *3.5*). As noted in the methods, the data used here was acquired across a period of time and a number of different sites, with specific acquisition parameters varying per participant. Such site and acquisition differences could potentially influence the derived metrics used in the current study. For the macrostructural measures, the hippocampal segmentations across all participants were quality controlled, ideally ensuring that regardless of site or structural acquisition, the segmentations were of sufficient quality for analysis. For the diffusion data, we found that for each group the extent of data acquired on the b=1 ms/μm^2^ was similar. A further limitation of the diffusion data was that opposite phase-encoding data was not available to perform susceptibility-induced distortion correction. Though the diffusion data was registered to the anatomical image, which may help reduce some of the distortions.

## 5. Conclusion

Alzheimer’s disease is a debilitating neurodegenerative disease which is characterized by large structural changes including the presence of neurofibrillary tangles, accumulation of extracellular amyloid-beta, glial activation, neuronal loss and corresponding deficits in cognition including memory impairment. The hippocampus is a key orchestrator in both encoding and retrieval of memory, and as such has been a major target for many facets of AD research. In the current study we looked to examine the macro- and microstructure throughout the whole spatial extent of the hippocampus in AD, MCI, and healthy aged controls. We found distinct decreases in both thickness and gyrification in AD and MCI across the subfields and anterior-posterior axis.

Related to the microstructure, we found significant increases in mean diffusivity in AD and MCI, which could be related to loss of neuronal cells, decreased fiber density, demyelination, and increased presence of CSF. We then examined the orientation of the primary direction of diffusion relative to the three canonical hippocampal axes and found distinct diffusion orientation shifts in AD and MCI throughout the anterior-posterior extent of the subiculum and CA1. Such AD-related changes in diffusion orientations were found to not have significant spatial overlap with AD-related changes in MD, suggesting that they may be capturing distinct spatially-localized disease processes. Finally, we showed that the macro- and microstructure of the hippocampus in AD appeared to change less across age relative to MCI and controls, and that age-related hippocampal macrostructure changes in MCI are similar and perhaps not distinguishable from healthy aging. Overall, this work provided a detailed characterization of the hippocampus in AD and MCI, and showed the potential utility of a new measure related to the orientation of diffusion.

## Declaration of Competing Interests

The authors have no competing interests to declare.

## Data and code availability

The data from this study was obtained from the Alzheimer’s Disease Neuroimaging Initiative (ADNI) database. The code used for the calculation of the cosine similarities can be found at: https://github.com/Bradley-Karat/Hippo_Macro_Micro

## Author Contributions

**Bradley G. Karat:** Conceptualization, Methodology, Software, Validation, Formal analysis, Writing - Original Draft, Writing - Review & Editing, Visualization

**Mahdieh Varvani Farahani:** Validation, Writing - Review & Editing

**Malcolm Davidson:** Validation, Writing - Review & Editing

**Arun Thurairajah:** Validation, Writing - Review & Editing

**Alaa Taha:** Validation, Writing - Review & Editing

**Taylor W. Schmitz:** Formal analysis, Writing - Review & Editing

**Ali R. Khan:** Conceptualization, Formal analysis, Software, Resources, Supervision, Writing - Review & Editing, Visualization, Funding acquisition

## Funding

BGK was supported by a post-graduate scholarship from the Natural Sciences and Engineering Research Council of Canada (NSERC). ARK was supported by the Canada Research Chairs program #950-231964, NSERC Discovery Grant RGPIN-2023-05558, Canada Foundation for Innovation (CFI) John R. Evans Leaders Fund project #37427, the Canada First Research Excellence Fund, and Brain Canada.

## Acknowledgements

Data collection and sharing for the Alzheimer’s Disease Neuroimaging Initiative (ADNI) is funded by the National Institute on Aging (National Institutes of Health Grant U19AG024904). The grantee organization is the Northern California Institute for Research and Education. In the past, ADNI has also received funding from the National Institute of Biomedical Imaging and Bioengineering, the Canadian Institutes of Health Research, and private sector contributions through the Foundation for the National Institutes of Health (FNIH) including generous contributions from the following: AbbVie, Alzheimer’s Association; Alzheimer’s Drug Discovery Foundation; Araclon Biotech; BioClinica, Inc.; Biogen; BristolMyers Squibb Company; CereSpir, Inc.; Cogstate; Eisai Inc.; Elan Pharmaceuticals, Inc.; Eli Lilly and Company; EuroImmun; F. Hoffmann-La Roche Ltd and its affiliated company Genentech, Inc.; Fujirebio; GE Healthcare; IXICO Ltd.; Janssen Alzheimer Immunotherapy Research & Development, LLC.; Johnson & Johnson Pharmaceutical Research & Development LLC.; Lumosity; Lundbeck; Merck & Co., Inc.; Meso Scale Diagnostics, LLC.; NeuroRx Research; Neurotrack Technologies; Novartis Pharmaceuticals Corporation; Pfizer Inc.; Piramal Imaging; Servier; Takeda Pharmaceutical Company; and Transition Therapeutics

## Notes

### Competing Interest Statement

The authors have declared no competing interest.

