## Supplementary figures for "Hippocampal grey matter changes across scales in Alzheimer’s Disease"

### Supplementary material

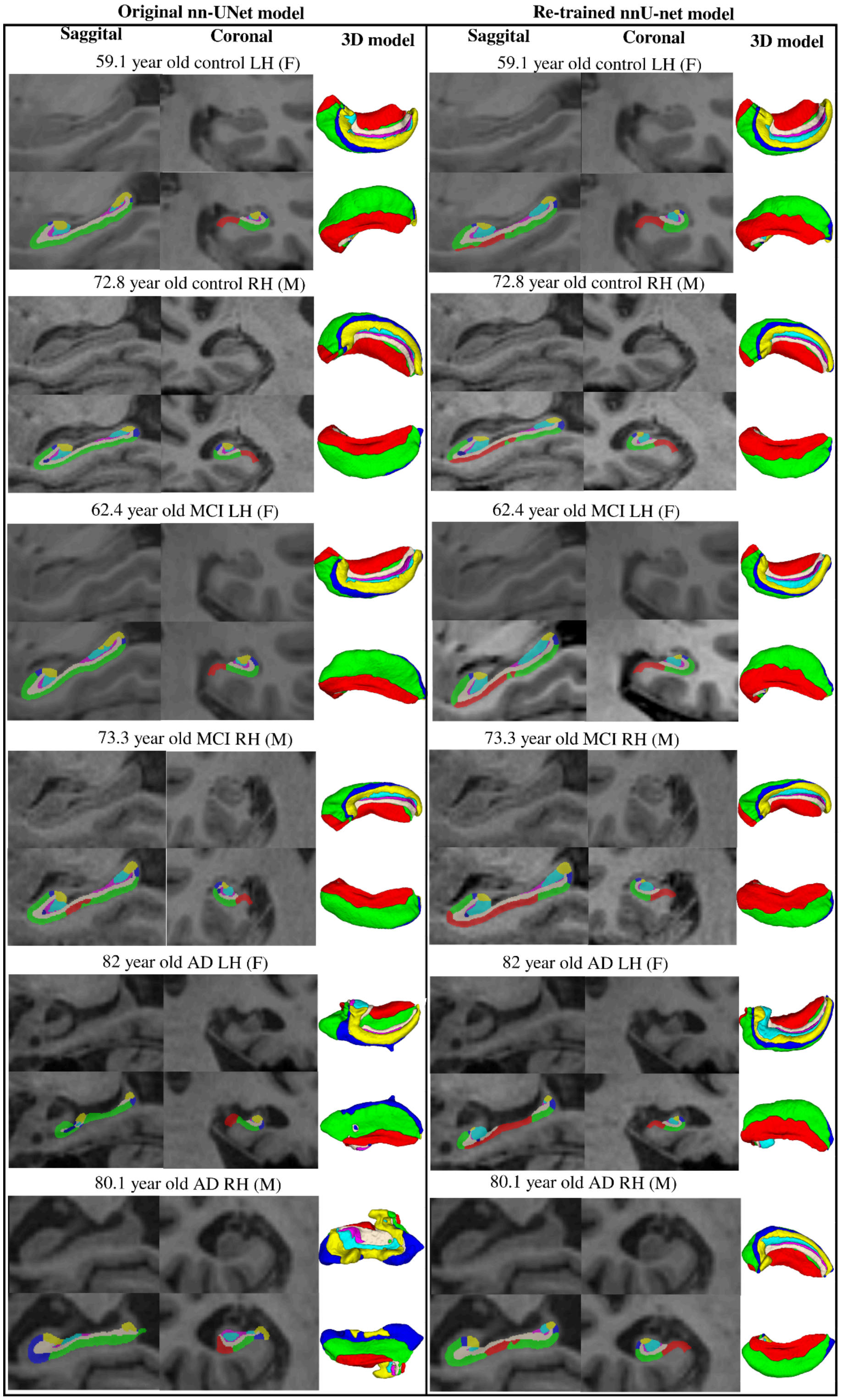


**Supplementary Figure 1.** Examples of HippUnfold segmentations with the original (left) and re-trained (right) nnU-net models across a handful of controls, MCI, and AD participants.

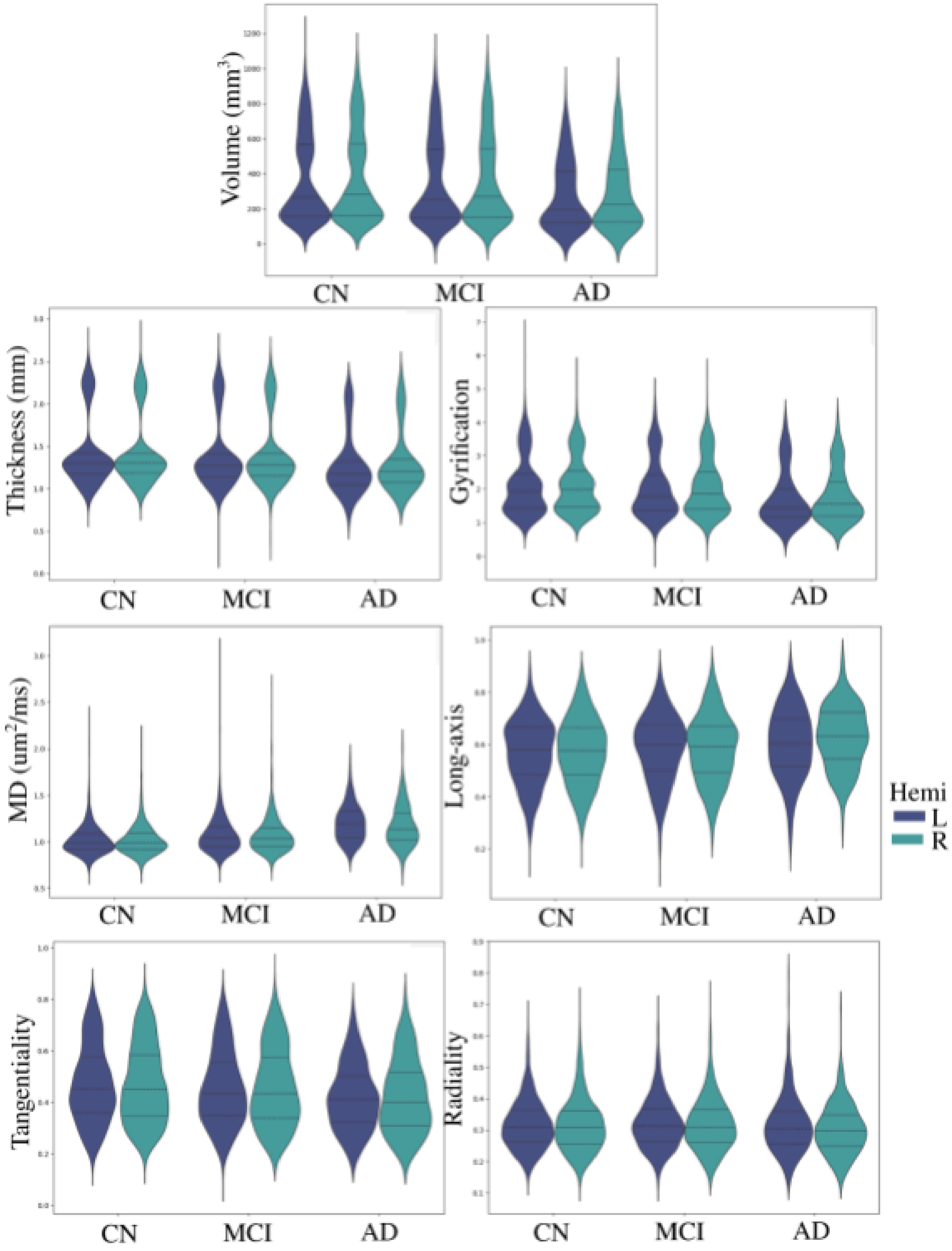


**Supplementary Figure 2.** Depicting all macro- and microstructural measures within each group separately for the left and right hemisphere. The lines within each distribution represent the quartiles of the data.

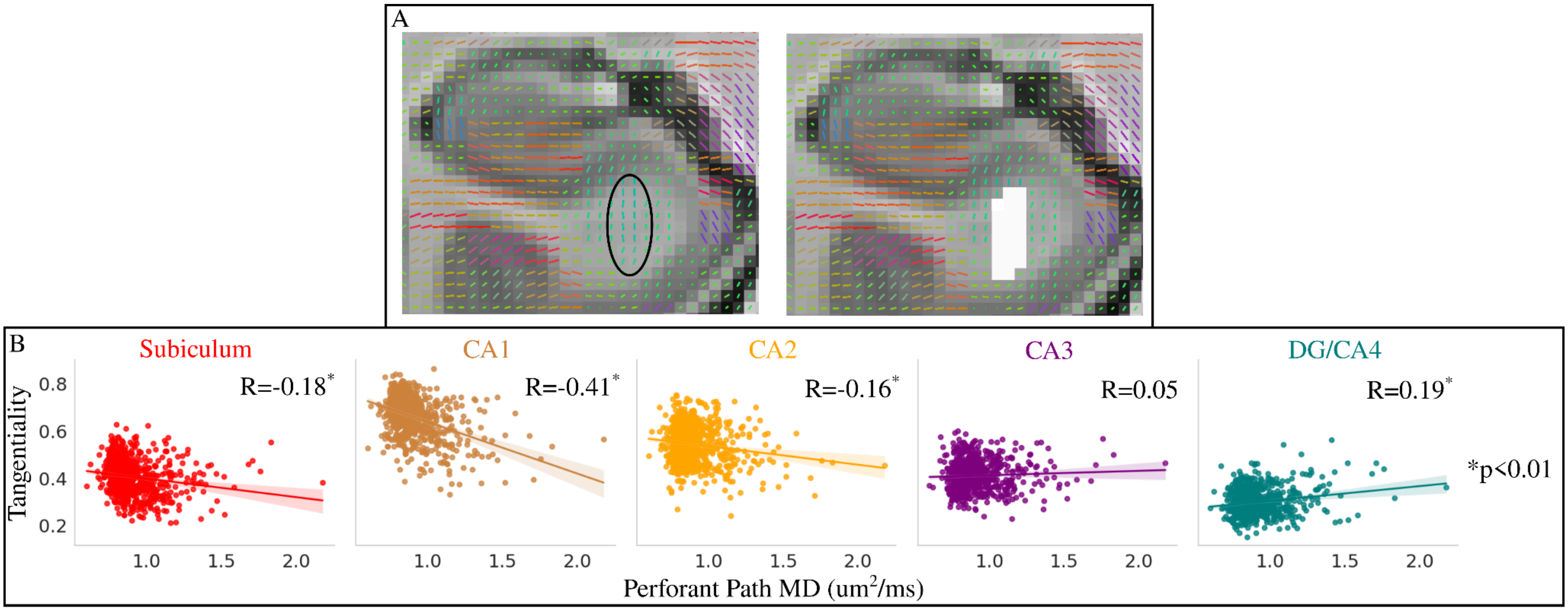


**Supplementary Figure 3.** Correlation between the mean diffusivity (MD) of a coarse perforant path (PP) segmentation and subfield-averaged tangentiality. (A) Coronal slice of a T1w image with V1 overlaid. The approximate location of the PP was determined by the orientation of V1 between the entorhinal cortex and the subiculum (Yassa et al., 2010). Black ellipsoid depicts the voxels which have orientations which align with the known trajectory of PP fibers. In the top right the white depicts the segmentation of the PP for that slice. This method was repeated on coronal slices throughout the anterior and body of the hippocampus. (B) Correlation between PP MD and subfield-averaged tangentiality, quantified with Pearson's R. An asterisk represents a significant Pearson's R, with a Bonferroni correction for 5 tests ($p<0.01$).

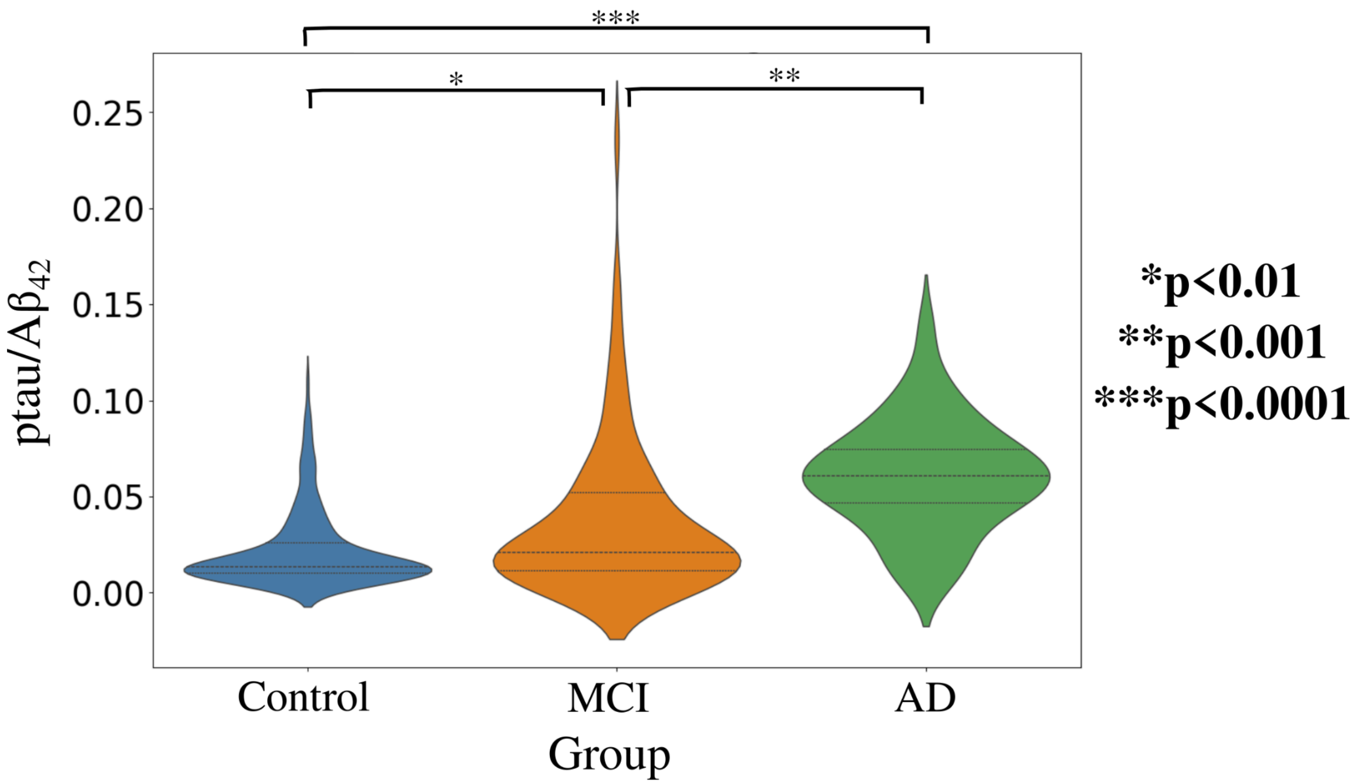


**Supplementary Figure 4**. Distributions of the ratio of phosphorylated tau (ptau) and $A\beta_{42}$ in a subset of 348 participants grouped by their diagnostic label determined through cognitive testing. Significant differences between groups are represented by asterisks, calculated with a Games-Howell post-hoc test after a significant Welch's ANOVA.

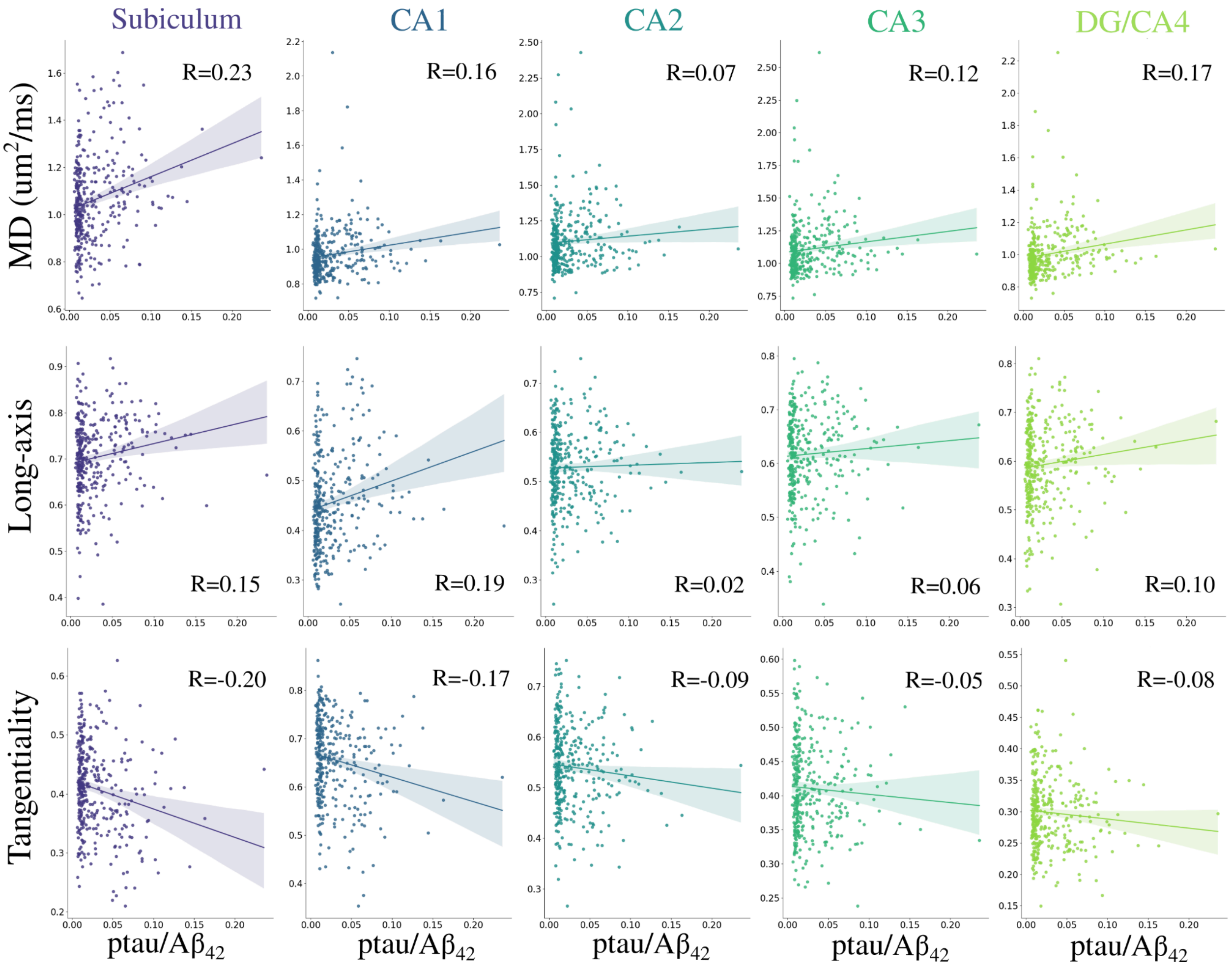


**Supplementary Figure 5.** Correlation between the subfield-averaged mean diffusivity (MD), long-axis, and tangential oriented diffusion with the ratio of phosphorylated tau (ptau) and $A\beta_{42}$ quantified using Pearson's R.

**Supplementary Table 1.** Differences in macro- and microstructural measures between controls (CN), mild cognitive impairment (MCI) and Alzheimer's disease (AD) for each subfield using the pairwise Games-Howell post-hoc test after a significant Welch's ANOVA.

| **Volume** | | | | | **Volume** | | | | |
|---|---|---|---|---|---|---|---|---|---|
| **Subiculum** | Comparison (A vs B) | M(A); M(B) | SE | *t* (df) | **CA1** | Comparison (A vs B) | M(A); M(B) | SE | *t* (df) |
| | AD vs CN | 405.5; 534.9 | 9.7 | -13.4(100.5)$^{***}$ | | AD vs CN | 606.4; 782.1 | 15.5 | -11.4(97.3)$^{***}$ |
| | AD vs MCI | 405.5; 500.6 | 10.8 | -8.8(149.6)$^{***}$ | | AD vs MCI | 606.4; 739.9 | 16.6 | -8.1(125.2)$^{***}$ |
| | CN vs MCI | 534.9; 500.6 | 6.8 | 5.1(381.5)$^{***}$ | | CN vs MCI | 782.1; 839.9 | 9.3 | 4.6(423.0)$^{***}$ |
| **Volume** | | | | | **Volume** | | | | |
| **CA2** | Comparison (A vs B) | M(A); M(B) | SE | *t* (df) | **CA3** | Comparison (A vs B) | M(A); M(B) | SE | *t* (df) |
| | AD vs CN | 115.3; 137.3 | 3.4 | -6.4(92.6)$^{***}$ | | AD vs CN | 210.8; 274.5 | 6.5 | -9.8(91.9)$^{***}$ |
| | AD vs MCI | 115.3; 133.2 | 3.6 | -5.0(107.4)$^{***}$ | | AD vs MCI | 210.8; 256.7 | 6.8 | -6.7(112.7)$^{***}$ |
| | CN vs MCI | 534.9; 500.6 | 1.7 | 2.4(457.2) | | CN vs MCI | 274.5; 256.7 | 3.4 | 5.2(419.4)$^{***}$ |
| **Volume** | | | | | **Gyrification** | | | | |
| **DG/CA4** | Comparison (A vs B) | M(A); M(B) | SE | *t* (df) | **Subiculum** | Comparison (A vs B) | M(A); M(B) | SE | *t* (df) |
| | AD vs CN | 119.84; 169.8 | 4.1 | -12.3(95.1)$^{***}$ | | AD vs CN | 1.5; 2.1 | 0.04 | -14.0(96.3)$^{***}$ |
| | AD vs MCI | 119.84; 157.4 | 4.4 | -8.5(126.1)$^{***}$ | | AD vs MCI | 1.5; 1.9 | 0.05 | -9.2(149.5)$^{***}$ |
| | CN vs MCI | 169.76; 157.4 | 2.4 | 5.1(400.5)$^{***}$ | | CN vs MCI | 2.1; 1.9 | 0.03 | 5.0(358.9)$^{***}$ |
| **Gyrification** | | | | | **Gyrification** | | | | |
| **CA1** | Comparison (A vs B) | M(A); M(B) | SE | *t* (df) | **CA2** | Comparison (A vs B) | M(A); M(B) | SE | *t* (df) |
| | AD vs CN | 1.9; 2.3 | 0.04 | -8.9(95.7)$^{***}$ | | AD vs CN | 1.1; 1.2 | 0.03 | -4.3(94.4)$^{**}$ |
| | AD vs MCI | 1.9; 2.2 | 0.04 | -6.5(119.8)$^{***}$ | | AD vs MCI | 1.1; 1.2 | 0.03 | -3.4(116.3)$^{*}$ |
| | CN vs MCI | 2.3; 2.2 | 0.02 | 3.6(429.3)$^{**}$ | | CN vs MCI | 1.2; 1.2 | 0.01 | 1.3(431.4) |
| **Gyrification** | | | | | **Gyrification** | | | | |
| **CA3** | Comparison (A vs B) | M(A); M(B) | SE | *t* (df) | **DG/CA4** | Comparison (A vs B) | M(A); M(B) | SE | *t* (df) |
| | AD vs CN | 1.3; 1.5 | 0.02 | -8.9(91.8)$^{***}$ | | AD vs CN | 3.1; 3.5 | 0.06 | -6.7(98.7)$^{***}$ |
| | AD vs MCI | 1.3; 1.5 | 0.03 | -6.4(109.0)$^{***}$ | | AD vs MCI | 3.1; 3.4 | 0.06 | –5.1(112.6)$^{***}$ |
| | CN vs MCI | 1.5; 1.5 | 0.01 | 4.5(438.3)$^{***}$ | | CN vs MCI | 3.5; 3.4 | 0.03 | 2.5(493.9) |

| | | | | | | | | | |
|---|---|---|---|---|---|---|---|---|---|
| **Thickness** | | | | | **Thickness** | | | | |
| **Subiculum** | Comparison (A vs B) | M(A); M(B) | SE | *t* (df) | **CA1** | Comparison (A vs B) | M(A); M(B) | SE | *t* (df) |
| | AD vs CN | 1.2; 1.3 | 0.01 | -9.0(96.4)$^{***}$ | | AD vs CN | 1.3; 1.4 | 0.01 | -8.4(94.7)$^{***}$ |
| | AD vs MCI | 1.2; 1.2 | 0.01 | -5.8(136.2)$^{***}$ | | AD vs MCI | 1.3; 1.3 | 0.01 | -5.9(115.2)$^{***}$ |
| | CN vs MCI | 1.3; 1.2 | 0.01 | 4.1(382.9)$^{**}$ | | CN vs MCI | 1.4; 1.3 | 0.01 | 4.0(439.1)$^{**}$ |
| **Thickness** | | | | | **Thickness** | | | | |
| **CA2** | Comparison (A vs B) | M(A); M(B) | SE | *t* (df) | **CA3** | Comparison (A vs B) | M(A); M(B) | SE | *t* (df) |
| | AD vs CN | 1.0; 1.1 | 0.01 | -3.6(91.5)$^{*}$ | | AD vs CN | 1.1; 1.3 | 0.02 | -9.8(88.1)$^{***}$ |
| | AD vs MCI | 1.0; 1.1 | 0.01 | -3.1(103.8)$^{*}$ | | AD vs MCI | 1.1; 1.2 | 0.02 | –6.9(105.9)$^{***}$ |
| | CN vs MCI | 1.1; 1.1 | 0.01 | 0.9(468.3) | | CN vs MCI | 1.3; 1.2 | 0.01 | 5.6(403.1)$^{***}$ |
| **Thickness** | | | | | **MD** | | | | |
| **DG/CA4** | Comparison (A vs B) | M(A); M(B) | SE | *t* (df) | **Subiculum** | Comparison (A vs B) | M(A); M(B) | SE | *t* (df) |
| | AD vs CN | 2.0; 2.2 | 0.03 | -9.5(85.9)$^{***}$ | | AD vs CN | 1.3; 1.0 | 0.03 | 8.8(95.6)$^{***}$ |
| | AD vs MCI | 2.0; 2.2 | 0.03 | -6.8(103.2)$^{***}$ | | AD vs MCI | 1.3; 1.1 | 0.03 | 6.1(116.0)$^{***}$ |
| | CN vs MCI | 2.2; 2.2 | 0.01 | 5.4(380.5)$^{***}$ | | CN vs MCI | 1.0; 1.1 | 0.01 | -4.3(445.0)$^{***}$ |
| **MD** | | | | | **MD** | | | | |
| **CA1** | Comparison (A vs B) | M(A); M(B) | SE | *t* (df) | **CA2** | Comparison (A vs B) | M(A); M(B) | SE | *t* (df) |
| | AD vs CN | 1.1; 1.0 | 0.02 | 6.6(90.3)$^{***}$ | | AD vs CN | 1.2; 1.1 | 0.03 | 6.2(95.7)$^{***}$ |
| | AD vs MCI | 1.1; 1.0 | 0.02 | 4.8(109.7)$^{***}$ | | AD vs MCI | 1.2; 1.1 | 0.03 | 4.1(127.0)$^{**}$ |
| | CN vs MCI | 1.0; 1.0 | 0.01 | -3.2(414.8)$^{*}$ | | CN vs MCI | 1.1; 1.1 | 0.02 | -2.9(402.8) |
| **MD** | | | | | **MD** | | | | |
| **CA3** | Comparison (A vs B) | M(A); M(B) | SE | *t* (df) | **DG/CA4** | Comparison (A vs B) | M(A); M(B) | SE | *t* (df) |
| | AD vs CN | 1.3; 1.1 | 0.02 | 7.9(99.3)$^{***}$ | | AD vs CN | 1.2; 1.0 | 0.02 | 10.0(93.2)$^{***}$ |
| | AD vs MCI | 1.3; 1.1 | 0.03 | 5.2(140.4)$^{***}$ | | AD vs MCI | 1.2; 1.0 | 0.02 | 6.1(138.6)$^{***}$ |
| | CN vs MCI | 1.1; 1.1 | 0.02 | -3.3(394.1)$^{*}$ | | CN vs MCI | 1.0; 1.0 | 0.01 | -5.0(353.5)$^{***}$ |
| **Long-axis** | | | | | **Long-axis** | | | | |
| **Subiculum** | Comparison (A vs B) | M(A); M(B) | SE | *t* (df) | **CA1** | Comparison (A vs B) | M(A); M(B) | SE | *t* (df) |
| | AD vs CN | 0.75; 0.69 | 0.01 | 6.4(121.2)$^{***}$ | | AD vs CN | 0.54; 0.44 | 0.01 | 9.0(96.7)$^{***}$ |

| | | | | | | | | | |
|---|---|---|---|---|---|---|---|---|---|
| | AD vs MCI | 0.75; 0.72 | 0.01 | 3.3(171.1)* | | AD vs MCI | 0.54; 0.47 | 0.01 | 6.2(117.4)*** |
| | CN vs MCI | 0.69; 0.72 | 0.01 | -3.4(439.6)* | | CN vs MCI | 0.44; 0.47 | 0.01 | -4.4(449.0)*** |
| **Tangential** | | | | | **Tangential** | | | | |
| **Subiculum** | Comparison (A vs B) | M(A); M(B) | SE | *t* (df) | **CA1** | Comparison (A vs B) | M(A); M(B) | SE | *t* (df) |
| | AD vs CN | 0.35; 0.42 | 0.01 | -7.9(102.5)*** | | AD vs CN | 0.58; 0.67 | 0.01 | -8.7(97.5)*** |
| | AD vs MCI | 0.35; 0.40 | 0.01 | -4.2(134.5)** | | AD vs MCI | 0.58; 0.65 | 0.01 | -6.2(127.6)*** |
| | CN vs MCI | 0.42; 0.40 | 0.01 | 5.3(433.1)*** | | CN vs MCI | 0.67; 0.65 | 0.01 | 3.4(416.8)* |
| **Radial** | | | | | **Radial** | | | | |
| **Subiculum** | Comparison (A vs B) | M(A); M(B) | SE | *t* (df) | **CA2** | Comparison (A vs B) | M(A); M(B) | SE | *t* (df) |
| | AD vs CN | 0.25; 0.28 | 0.01 | -3.8(126.0)** | | AD vs CN | 0.32; 0.29 | 0.01 | 2.7(100.4) |
| | AD vs MCI | 0.25; 0.28 | 0.01 | -3.2(164.7)* | | AD vs MCI | 0.32; 0.30 | 0.01 | 1.4(116.9) |
| | CN vs MCI | 0.28; 0.28 | 0.01 | 0.4(472.8) | | CN vs MCI | 0.29; 0.30 | 0.01 | -2.1(485.9) |

*p<0.01;**p<0.001; ***p<0.0001. SE - Standard error. M - Mean. df - Degrees of Freedom

**Supplementary Table 2.** Differences in macro- and microstructural measures between controls (CN), mild cognitive impairment (MCI) and Alzheimer's disease (AD) across the anterior-posterior axis using the pairwise Games-Howell post-hoc test after a significant Welch's ANOVA.

| **Gyrification** | | | | | **Gyrification** | | | | |
|---|---|---|---|---|---|---|---|---|---|
| **Head** | Comparison (A vs B) | M(A); M(B) | SE | *t* (df) | **Body** | Comparison (A vs B) | M(A); M(B) | SE | *t* (df) |
| | AD vs CN | 2.0; 2.4 | 0.05 | -8.9(92.2)*** | | AD vs CN | 1.6; 2.0 | 0.03 | -13.9(98.5)*** |
| | AD vs MCI | 2.0; 2.3 | 0.05 | -6.5(116.7)*** | | AD vs MCI | 1.6; 1.9 | 0.03 | -9.5(135.7)*** |
| | CN vs MCI | 2.4; 2.3 | 0.03 | 3.6(405.0)** | | CN vs MCI | 2.0; 1.9 | 0.02 | 5.5(400.1)*** |
| **Gyrification** | | | | | **Thickness** | | | | |
| **Tail** | Comparison (A vs B) | M(A); M(B) | SE | *t* (df) | **Head** | Comparison (A vs B) | M(A); M(B) | SE | *t* (df) |
| | AD vs CN | 0.5; 0.7 | 0.02 | -13.1(93.7)*** | | AD vs CN | 1.3; 1.4 | 0.01 | -9.1(88.0)*** |
| | AD vs MCI | 0.5; 0.7 | 0.02 | -9.2(121.6)*** | | AD vs MCI | 1.3; 1.3 | 0.01 | -6.2(111.9)*** |
| | CN vs MCI | 0.7; 0.7 | 0.01 | 5.7(403.1)*** | | CN vs MCI | 1.4; 1.3 | 0.01 | 4.9(373.2)*** |
| **Thickness** | | | | | **Thickness** | | | | |

| **Body** | Comparison (A vs B) | M(A); M(B) | SE | *t* (df) | **Tail** | Comparison (A vs B) | M(A); M(B) | SE | *t* (df) |
|---|---|---|---|---|---|---|---|---|---|
| | AD vs CN | 1.3; 1.3 | 0.01 | -8.3(93.8)$^{***}$ | | AD vs CN | 1.2; 1.4 | 0.02 | -10.8(86.0)$^{***}$ |
| | AD vs MCI | 1.3; 1.3 | 0.01 | -5.8(117.3)$^{***}$ | | AD vs MCI | 1.2; 1.3 | 0.02 | -8.1(103.1)$^{***}$ |
| | CN vs MCI | 1.3; 1.3 | 0.01 | 4.0(421.0)$^{**}$ | | CN vs MCI | 1.4; 1.3 | 0.01 | 5.2(383.6)$^{***}$ |
| **MD** | | | | | **MD** | | | | |
| **Head** | Comparison (A vs B) | M(A); M(B) | SE | *t* (df) | **Body** | Comparison (A vs B) | M(A); M(B) | SE | *t* (df) |
| | AD vs CN | 1.2; 1.0 | 0.03 | 7.8(85.8)$^{***}$ | | AD vs CN | 1.2; 1.0 | 0.02 | 8.0(100.2)$^{***}$ |
| | AD vs MCI | 1.2; 1.0 | 0.03 | 5.5(100.8)$^{***}$ | | AD vs MCI | 1.2; 1.1 | 0.02 | 5.4(130.4)$^{***}$ |
| | CN vs MCI | 1.0; 1.0 | 0.01 | -4.9(394.9)$^{***}$ | | CN vs MCI | 1.0; 1.1 | 0.01 | -3.3(429.3)$^{*}$ |
| **MD** | | | | | **Long-axis** | | | | |
| **Tail** | Comparison (A vs B) | M(A); M(B) | SE | *t* (df) | **Head** | Comparison (A vs B) | M(A); M(B) | SE | *t* (df) |
| | AD vs CN | 1.3; 1.1 | 0.04 | 7.4(92.2)$^{***}$ | | AD vs CN | 0.51; 0.46 | 0.01 | 5.9(97.4)$^{***}$ |
| | AD vs MCI | 1.3; 1.1 | 0.04 | 5.1(111.2)$^{***}$ | | AD vs MCI | 0.51; 0.48 | 0.01 | 3.6(117.7)$^{**}$ |
| | CN vs MCI | 1.1; 1.1 | 0.02 | -4.0(429.2)$^{**}$ | | CN vs MCI | 0.46; 0.48 | 0.01 | -3.7(455.6)$^{**}$ |
| **Long-axis** | | | | | **Long-axis** | | | | |
| **Body** | Comparison (A vs B) | M(A); M(B) | SE | *t* (df) | **Tail** | Comparison (A vs B) | M(A); M(B) | SE | *t* (df) |
| | AD vs CN | 0.71; 0.63 | 0.01 | 6.9(102.6)$^{***}$ | | AD vs CN | 0.65; 0.57 | 0.01 | 5.7(109.1)$^{***}$ |
| | AD vs MCI | 0.71; 0.66 | 0.01 | 4.6(127.8)$^{***}$ | | AD vs MCI | 0.65; 0.61 | 0.01 | 2.6(141.9) |
| | CN vs MCI | 0.63; 0.66 | 0.01 | -3.2(456.4)$^{*}$ | | CN vs MCI | 0.57; 0.61 | 0.01 | -4.3(451.1)$^{***}$ |
| **Tangential** | | | | | **Tangential** | | | | |
| **Head** | Comparison (A vs B) | M(A); M(B) | SE | *t* (df) | **Body** | Comparison (A vs B) | M(A); M(B) | SE | *t* (df) |
| | AD vs CN | 0.56; 0.63 | 0.01 | -7.7(97.9)$^{***}$ | | AD vs CN | 0.39; 0.45 | 0.01 | -7.0(98.7)$^{***}$ |
| | AD vs MCI | 0.56; 0.61 | 0.01 | -5.2(133.5)$^{***}$ | | AD vs MCI | 0.39; 0.43 | 0.01 | -4.6(117.6)$^{***}$ |
| | CN vs MCI | 0.63; 0.61 | 0.01 | 3.3(401.5)$^{*}$ | | CN vs MCI | 0.45; 0.43 | 0.01 | 3.8(468.3)$^{**}$ |
| **Tangential** | | | | | **Radial** | | | | |
| **Tail** | Comparison (A vs B) | M(A); M(B) | SE | *t* (df) | **Head** | Comparison (A vs B) | M(A); M(B) | SE | *t* (df) |
| | AD vs CN | 0.37; 0.43 | 0.01 | -6.8(109.5)$^{***}$ | | AD vs CN | 0.34; 0.32 | 0.01 | 2.4(100.2) |
| | AD vs MCI | 0.37; 0.41 | 0.01 | -3.9(134.8)$^{*}$ | | AD vs MCI | 0.34; 0.33 | 0.01 | 1.1(133.8) |

| | | | | | | | | | |
|---|---|---|---|---|---|---|---|---|---|
| | CN vs MCI | 0.43; 0.41 | 0.01 | 4.0(476.1)** | | CN vs MCI | 0.32; 0.33 | 0.01 | -2.0(418.6) |

*p<0.017;**p<0.0017; ***p<0.00017. SE - Standard error. M - Mean. df - Degrees of Freedom
